# Neuronal calcium spikes enable vector inversion in the *Drosophila* brain

**DOI:** 10.1101/2023.11.24.568537

**Authors:** Itzel G. Ishida, Sachin Sethi, Thomas L. Mohren, Mia Haraguchi, L.F. Abbott, Gaby Maimon

**Author notes:** Lead Contact: Gaby Maimon (G.M.). Correspondence (G.M.).

## Abstract

A typical neuron signals to downstream cells when it is depolarized and firing sodium spikes. Some neurons, however, also fire calcium spikes when hyperpolarized. The function of such bidirectional signaling remains unclear in most circuits. Here we show how a neuron class that participates in vector computation in the fly central complex employs hyperpolarization-elicited calcium spikes to invert two-dimensional mathematical vectors. By switching from firing sodium to calcium spikes, these neurons implement a ∼180° realignment between the vector encoded in the neuronal population and the fly’s internal compass signal, thus inverting the vector. We show that the calcium spikes rely on the T-type calcium channel Ca-α1T, and argue, via analytical and experimental approaches, that these spikes enable vector computations in portions of angular space that would otherwise be inaccessible. These results reveal a seamless interaction between molecular, cellular and circuit properties for implementing vector mathematics in the brain.

## Introduction

Sensory processing takes place in *egocentric* coordinates. We see a visual stimulus on our left; we feel air blowing on the front of our face, and so on. In contrast, brain centers that compute navigation-related variables typically do so in map-like or *allocentric* coordinates^1^. We orient north on a hike; wind arrives from the east, and so on. Animals likely make use of such allocentric brain signals when performing feats such as migrating thousands of miles to breeding sites^2–6^. These signals also enable more humble tasks, like allowing fruit flies to maintain a consistent traveling direction for an hour when they wish to disperse far from a start location^7–9^, or allowing flies to remember a direction of interest for minutes after external cues informative of that direction have disappeared^10^. Because allocentric navigational signals are constructed from egocentric sensory inputs, brains must perform egocentric-to-allocentric coordinate transformations to build them^1^. Until recently, how such coordinate transformations are implemented was unclear outside of computational modeling. However, work in the *Drosophila* central complex has provided a detailed, biological example of how a coordinate transformation is implemented in the central brain^11,12^. This circuit transforms a fly’s direction of travel from egocentric (e.g, walking leftward) to allocentric (e.g., walking north) coordinates^11,12^.

The allocentric traveling direction signal is constructed by summing the activity of four presynaptic neuronal populations, each of which can be modeled as encoding a two-dimensional mathematical vector^11^. The lengths of these neuronally encoded vectors are controlled by the fly’s egocentric traveling direction and their angles are controlled by the fly’s allocentric heading angle (a separately constructed signal). Summing the activity of the four neuronal populations is functionally equivalent to performing vector addition of the four vectors they encode, with the vector sum accurately tracking the fly’s allocentric traveling angle^11^. Formally, the fly’s traveling angle could have been calculated by summing only two vectors––akin to x and y components in Cartesian geometry––if each component vector would have been able to point in both the positive and negative direction along its respective axis. However, the vectors in the traveling-direction system have never been observed to perform such an inversion.

In this study, we show how a different pair of neuronal populations in the fly central complex can calculate the allocentric direction of airflow using only two component vectors, with each vector being able to point in both the positive and negative direction along its respective axis. The constituent neurons––PFNa cells––signal vectors along the positive direction by firing canonical sodium spikes upon depolarization and signal vectors along the negative direction by firing non-canonical calcium spikes upon hyperpolarization. This dual-spike mechanism circumvents the standard limitations of rectification, allowing the system to encode both a vector and its inverse. We show that the hyperpolarization-elicited calcium spikes are mediated by T-type calcium channels, which are known to produce analogous calcium spikes and burst firing in mammalian thalamocortical circuits^13,14^. We additionally describe a population of neurons––which are monosynaptically downstream of PFNa cells––that can sum the PFNa vectors. These read-out neurons can sum the sodium-spike encoded vectors through standard synaptic transmission. The calcium spikes, on the other hand, are at times associated with a strong postsynaptic response in the read-out neurons and at other times are associated with much weaker responses, suggesting that the T-type encoded vectors might function in processes (e.g., vector integration) that transcend fast synaptic transmission. These results––supported by a model rooted in detailed anatomy and physiology––reveal a mechanism by which neurons can implement invertible, two-dimensional vectors.

## Results

### How neuronal populations could encode invertible, two-dimensional vectors

Vector computations in the fly central complex (**Figure 1A**) are anchored to a common, internally generated sense of heading signaled by EPG cells^8,15^ (**Figure 1B**, top). EPG cells tile a central-complex structure called the ellipsoid body with their dendrites, where they express a spatially localized calcium signal, or *bump*. This bump rotates around the ellipsoid body as the fly turns, with its position around the ellipsoid body, or *phase*, tracking the fly’s heading. There are two copies of the ellipsoid body bump evident in the EPG axons in a second central complex structure, called the protocerebral bridge (**Figure 1B**, top). The phase of the two bumps across the bridge, and the one bump around the ellipsoid body, all indicate the same variable: the fly’s orientation within an environment.

**Figure 1.**
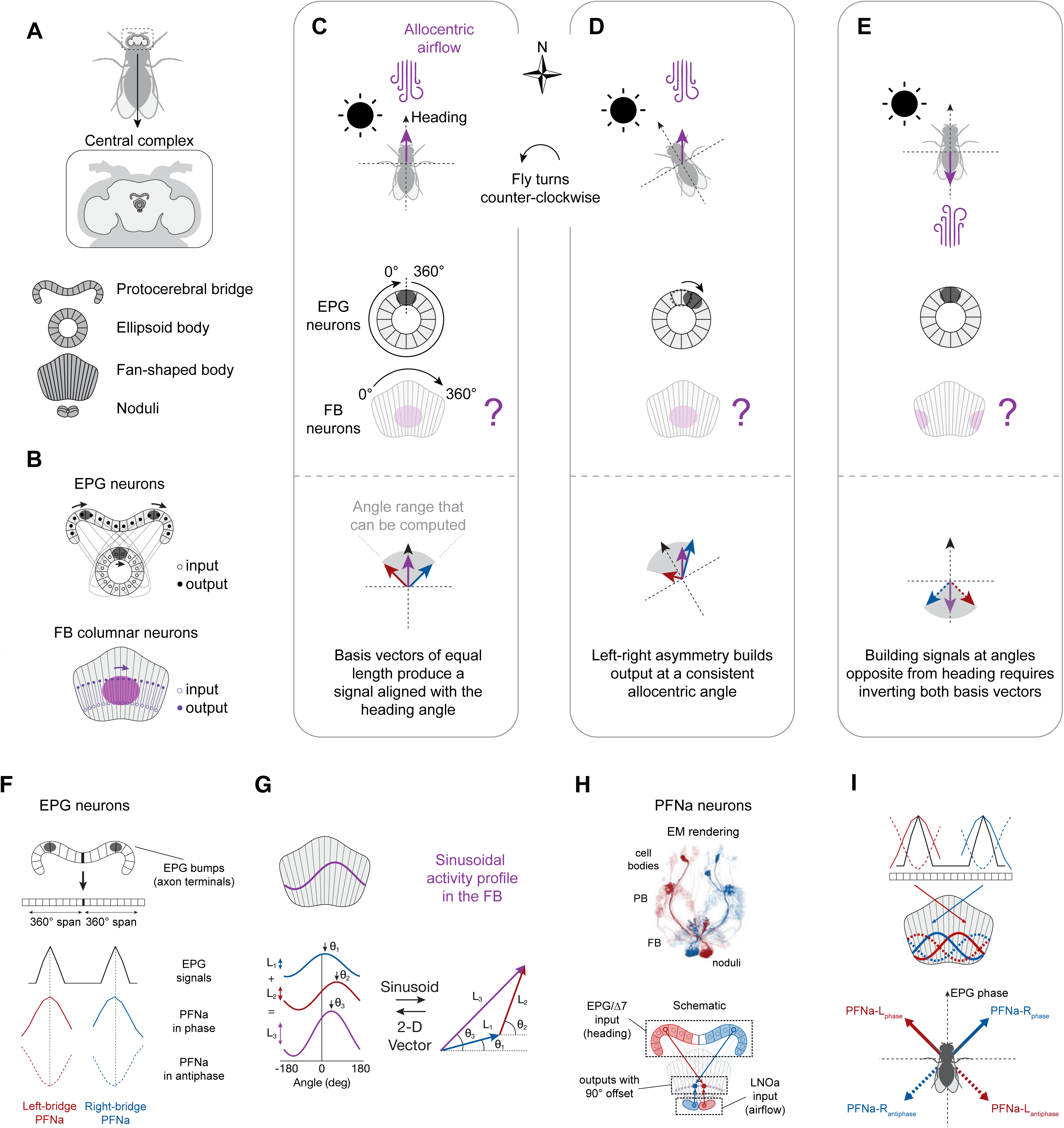
A framework for implementing coordinate transformations using invertible vectors in the *Drosophila* central complex. **(A)** *Drosophila* central complex. **(B)** EPG neurons and fan-shaped body neurons. **(C)** Top: A fly faces north while wind blows from the north. Middle: expected neuronal signals. Bottom: 2D vector expression. The red and blue arrows depict the basis vectors that could build an allocentric wind direction signal (purple). The gray region depicts the range of angles that could be computed by these basis vectors. **(D)** Same as in C, but the fly turned counterclockwise relative to C. **(E)** Same as in C, but with wind blowing from the south. **(F)** The two copies of the EPG signal are transmitted to the downstream PFNa cells. If the two PFNa activity peaks could flip 180°, this would be equivalent to inverting their encoded vector. **(G)** Sinusoidal profiles in the population activity of central complex neurons can be understood as vectors. The sinusoid’s amplitude is the 2D vector’s length; its phase is the 2D vector’s angle. **(H)** Electron microscopy reconstruction of the 58 PFNa neurons in the hemibrain connectome^17,60^. Cells that output onto the same neuron in the fan-shaped body are highlighted. The left- and right-bridge PFNa neurons project to the fan-shaped body with anatomical offsets that introduce ∼±45° rotations of their encoded vectors relative to the heading angle^17^. PFNa neurons that receive EPG input in the left bridge also receive input from the airflow-tuned LNOa neurons^21^ in the right nodulus, and vice versa. (I) The sinusoidal activity bumps in the PFNa neurons at the bridge are transmitted to the fan-shaped body with ∼±45° offsets relative to the EPG signal. If either PFNa sinusoids in the bridge were to invert their phase, then this would invert the direction of the encoded vectors to ∼±135° relative to the EPG signal.

Whereas the EPG phase tracks the fly’s heading, the position, or phase, of calcium bumps in a third central complex structure––the fan-shaped body––have been shown or hypothesized to signal additional variables in the EPG-yoked reference frame^10–12,16–20^ (**Figure 1B**, bottom). The fan-shaped body bumps are expressed by columnar populations of cells, each of which tiles the fan shaped body from left to right. We hypothesized that the phase of one of these fan-shaped body bumps signals the allocentric direction of airflow^21^. When a fly turns in the context of a constant wind direction, the EPG bump rotates in the ellipsoid body to indicate the new heading (**Figures 1C and 1D**, gray) but this putative fan-shaped body bump should remain stationary (**Figures 1C and 1D**, purple) because the allocentric airflow direction did not change. If the airflow direction flips 180°, the fan-shaped body bump should shift by half the width of the fan-shaped body, or 180° (**Figure 1E**). Does such an airflow signal exist in the fan-shaped body, and if so, how might it be built?

In the bridge, EPG cells provide synaptic input to dozens of downstream neurons, including several classes of PFN cells^17^. Whereas the two EPG calcium bumps in the bridge are often narrow in shape, the downstream pair of bumps in each class of PFN cells––one in the left bridge and one in the right bridge––are broader, conforming well to sinusoidal functions across the bridge^11,17^ (**Figure 1F**). The sinusoidal activity profile of each PFN population in the bridge appears to encode a two-dimensional vector in a so-called phasor representation (**Figure 1G**). The amplitude of the sinusoid encodes the vector’s length and its peak position, or phase, encodes the vector’s angle^11,16,22,23^. Adding two or more PFN sinusoids is equivalent to performing vector addition on the vectors they encode (**Figure 1G**).

The phases of the PFN bumps in the left and right bridge are yoked to the EPG phase, such that both PFN-encoded vectors point in the fly’s heading direction. When these PFNs project to the fan-shaped body, their projections are anatomically offset—rotating the right vector ∼45° clockwise and the left ∼45° counterclockwise—creating a ∼90° offset between the vector angles^11,17^ (**Figures 1H and 1I**). The length of each vector is scaled by input that the entire PFN population receives in the contralateral nodulus (a fourth central-complex structure); this nodulus input carries egocentric signals that are appropriate for determining vector length (**Figure 1H**). Finally, fan-shaped body neurons monosynaptically downstream of PFN cells add, column by column, the sinusoidal activity patterns across two or more PFN populations^11^, producing a sinusoidal activity pattern whose amplitude and phase match the length and angle of the vector sum of the input vectors.

Adding a pair of vectors with a 90° offset allows the output vector to point anywhere within a 90° sector of angular space (**Figures 1C and 1D**, bottom vector diagrams). Other parts of angular space can only be covered by including a second vector pair^11,12,16^ or by allowing the vectors within the one pair to invert their direction (**Figure 1E**, bottom vector diagram). We thus hypothesized that there might exist situations in which PFN-encoded vectors could invert, that is, flip from pointing toward the positive direction of their particular axis to pointing toward the negative direction. Physiologically, when a PFN vector points in the positive direction, one would observe the standard alignment between the peak of the PFN sinusoid and the EPG bump driving it in the bridge (**Figure 1I**, solid lines). When the vector points in the inverse direction, there would be a ∼180° offset between the PFN and EPG signals in the bridge (**Figure 1I**, dotted lines). A 180° offset of a phasor is equivalent to inverting the direction of the encoded vector.

## Population calcium signals in PFNa neurons signal invertible vectors

We hypothesized that PFNa cells––a specific PFN subtype that is responsive to airflow^21^––might signal two vectors that can invert. Most of the synaptic output of PFNa neurons is onto cells that do not receive input from other PFN neurons^17^, making it hard to imagine how a four-vector circuit motif, akin to the one employed by the traveling direction system^11,12^, could be implemented. In addition, as will be evident below, PFNa neurons do not show clear, heading-related, bumps of activity when a fly explores a virtual reality environment that lacks airflow stimuli, which could be due to the two vectors inverting their phase irregularly when their amplitudes are minimal in the context of no airflow. Most other columnar bridge neurons have a measurable pair of bumps in the bridge in the context of a visual virtual environment^11,12,24–26^.

To study PFNa activity, we placed head-fixed flies on an air-cushioned ball and allowed the flies to navigate in a simple virtual environment as we imaged neural activity with a two-photon microscope (**Figure 2A**). The flies’ left/right turns controlled the position of a bright vertical bar on a panoramic visual display, such that the bar’s angular position tracked the fly’s heading in the virtual world akin to a distant cue in the real world, like the sun. Because PFNa neurons have been previously shown to be responsive to airflow stimuli^21^, they seemed poised to transform the direction of airflow from egocentric to allocentric coordinates^21^. Thus, we also surrounded the flies with a ring of 36 static tubes that allowed us to deliver airflow stimuli from various directions (**Methods, Figure 2A**).

**Figure 2.**
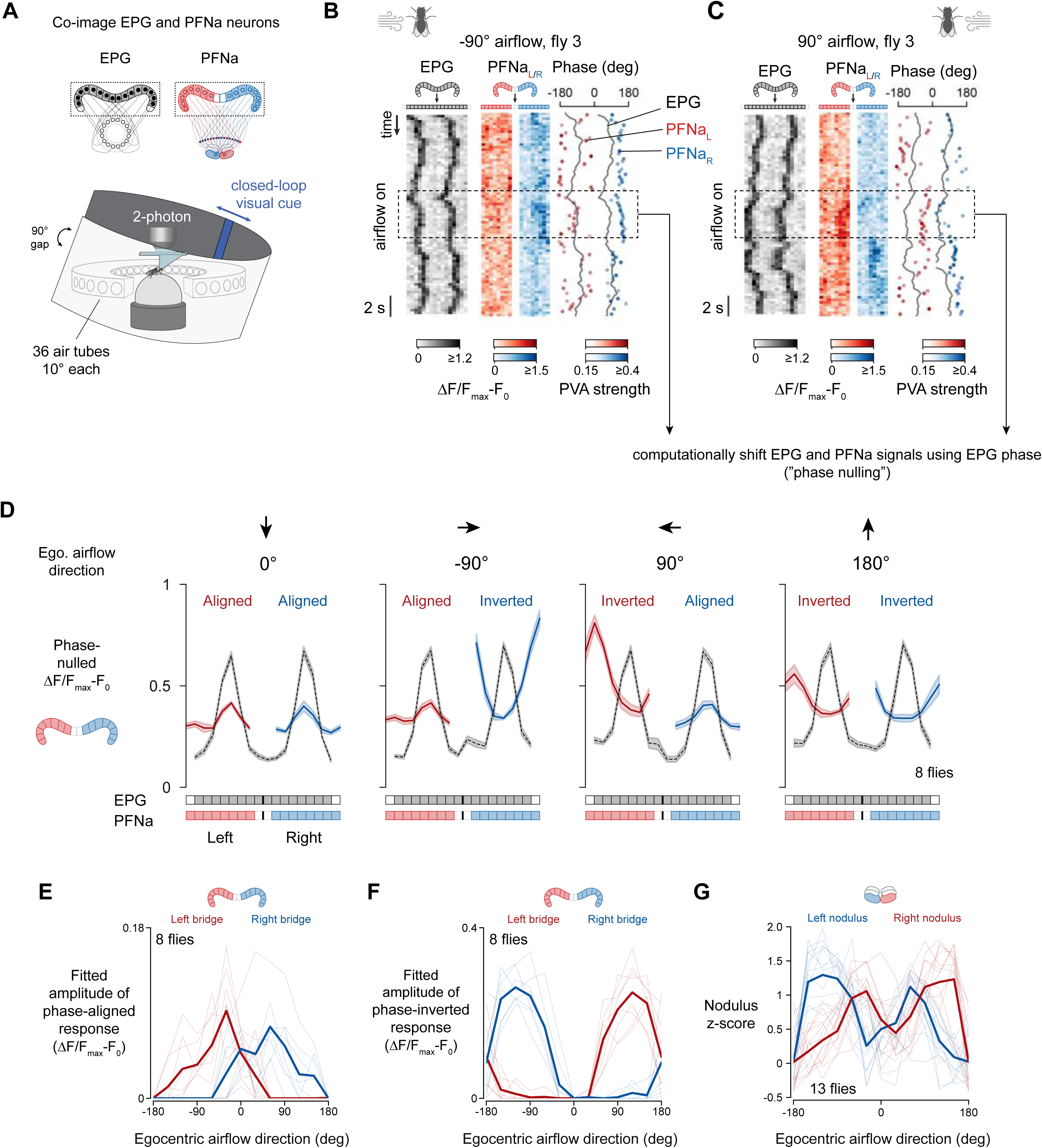
PFNa neurons express activity bumps that can be both in-phase and antiphase relative to the EPG bumps in the protocerebral bridge. **(A)** Imaging neural activity in a fly walking with a constant heading cue and intermittent air puffs. **(B and C)** Simultaneous calcium imaging of EPG and PFNa neurons in the protocerebral bridge. Example responses to an air puff from the fly’s left (-90°) or right (90°). Gray: jRGECO1a signal from EPG neurons. Red and blue: jGCaMP7f signal from PFNa neurons. Right: estimated phases of the EPG, left-bridge PFNa, and right-bridge PFNa bumps. Dot color shows the population vector average (PVA) strength. **(D)** Phase-nulled EPG and PFNa signals averaged over time. Mean and SEM shown. **(E and F)** Amplitude of the phase-aligned and phase-inverted bumps in the population activity of PFNa neurons, determined from a fit (Methods). **(G)** Mean PFNa activity in the noduli as a function of airflow direction.

One complication is that airflow can sometimes cause the EPG heading bump (i.e., the fly’s internal sense of heading) to rotate^27^. We verified that that the EPG phase did not rotate with the brief, randomly directed open-loop puffs used in these experiments (**Figure S1A-S1I**). In other words, the fly seemed to interpret each air puff as a disturbance arriving from a different allocentric direction rather than its body having turned in the context of a static wind direction. We could thus study how PFNa neurons signal the direction of each puff.

We co-imaged calcium in the axon terminals of EPG cells and in the dendrites of PFNa cells in the protocerebral bridge as flies walked with a closed-loop bar stimulus and experienced open-loop air puffs from various directions. The EPG bumps in the bridge were consistently measurable during these experiments. The PFNa bumps, on the other hand, were often dim, but typically became clearer during puffs (**Figures 2B and 2C**). Notably, when we puffed air on a single fly from the left, the peak of the left-bridge PFNa bump was aligned to the peak of the EPG signal, but the peak of the right-bridge PFNa bump was offset from the EPG peak by ∼180° (**Figure 2B**, dotted box). When we puffed air on the right side of the same fly, the opposite relationship was evident (**Figure 2C**, dotted box). These findings stand in contrast to what is observed in the traveling-direction system, where the activity bumps of all four PFN populations always align with the EPG peak in the bridge^11,12^.

To test whether the examples shown in Figure 2B-C represented a consistent feature of the data, we computationally shifted the EPG bumps to the same angular position in the bridge and rotated the PFNa bumps by this same EPG-determined angle^11,24^ (**Methods**). When we plotted the mean across all flies of these EPG-phase-aligned bumps (**Figures 2D and S2A**), we found that, as in the example fly, the PFNa bumps could exist both in phase and antiphase with the EPG heading signal, contingent on the egocentric airflow direction (**Figure 2D**). Moreover, whether the PFNa bumps were dominantly in phase or antiphase, we observed that they were consistently well fit by sinusoidal functions, consistent with the bumps encoding 2D vectors (**Figure S2I**). With air puffs from the front, both the left and the right PFNa bumps had their peaks aligned with the EPG peaks (**Figures 2D, 2E, and S2A**). With air from the side, the PFNa sinusoid contralateral to the stimulated side of the body had its phase offset by ∼180° relative to the EPG bump (**Figures 2D and S2A**). With air from behind, both PFNa sinusoids expressed an antiphase relationship to the EPG bump (**Figures 2D, 2F, S2A, S2G, and S2H**). Thus, both PFNa sinusoids in the bridge invert, or equivalently shift their phase by ∼180°, when air puffs arrive from the back and one or the other sinusoid inverts when air comes from the side.

The above logic explains the PFNa vector direction, but not length. Vector length corresponds to the amplitude of the PFN phasor in the bridge and it can also be extracted from the PFN calcium signals in the noduli^11^ (**Figure 1A and 1H**). All left-bridge PFNa cells innervate the right nodulus, and vice versa. In the nodulus, each PFN population receives inputs from neurons that are sensitive to the egocentric variable the scales the length of the vector^11,16^. In the case of PFNa cells, as airflow rotates around the fly, our vector-inversion model predicts two activity peaks in the nodulus, 180° apart ––once when the airflow angle aligns with the positive direction of each vector’s projection axis and once when the airflow angle aligns with the negative direction. We indeed observed two peaks in the PFNa calcium signal in the nodulus as a function of the airflow angle around the fly, for both the left- and right-bridge PFNa cells (**Figures 2G, S2B, and S2C**), supporting the inversion model.

In principle, PFNa cells might have inherited their dual-peaked airflow-direction tuning curves from the neurons that bring in airflow signals to the nodulus: LNOa cells^17,21^. However, we found that the airflow tuning curves of LNOa cells have only a single peak (**Figures S2D and S2E**). How is it that PFNa cells show two calcium response peaks in the nodulus as a function of the airflow direction, when their inputs should cause them to maximally depolarize at only one preferred angle? Moreover, what is the mechanism that allows the PFNa sinusoids in the bridge to become phase inverted relative to the EPG signal when airflow arrives from certain directions and not others? As we show next, electrophysiological measurements from PFNa cells provided clear answers to these questions.

### PFNa neurons exhibit two types of spikes poised to underlie signaling of phase-aligned and phase-inverted vectors

We performed whole-cell patch-clamp recordings from PFNa neurons in head-fixed flies navigating with a closed-loop bar while they also received air puffs from various directions (**Figures 3A and 3B**). Consistent with a past report^21^, we observed strong responses from PFNa cells to air puffs, both in their membrane potential (*Vm*) and in their spike rate (**Figure 3B**; note that sodium spikes in PFNa cells have a very small amplitude when recorded at the soma and are hardly visible at the scale used in **Figure 3B**). PFNa electrical activity was also strongly modulated by heading (**Figure 3B**). Across a population of 20 single-neuron recordings, we observed strong tuning at the level of the mean (spike removed) *Vm* to both the fly’s heading and air-puff direction (**Figures 3C, 3D, 3E, S3A, S3B, S3D, and S3E**). The *Vm* tuning curves for both variables conformed well to sinusoidal functions (**Figure S2J**), as would be expected from neurons that participate in encoding vectors via a phasor representation.

**Figure 3.**
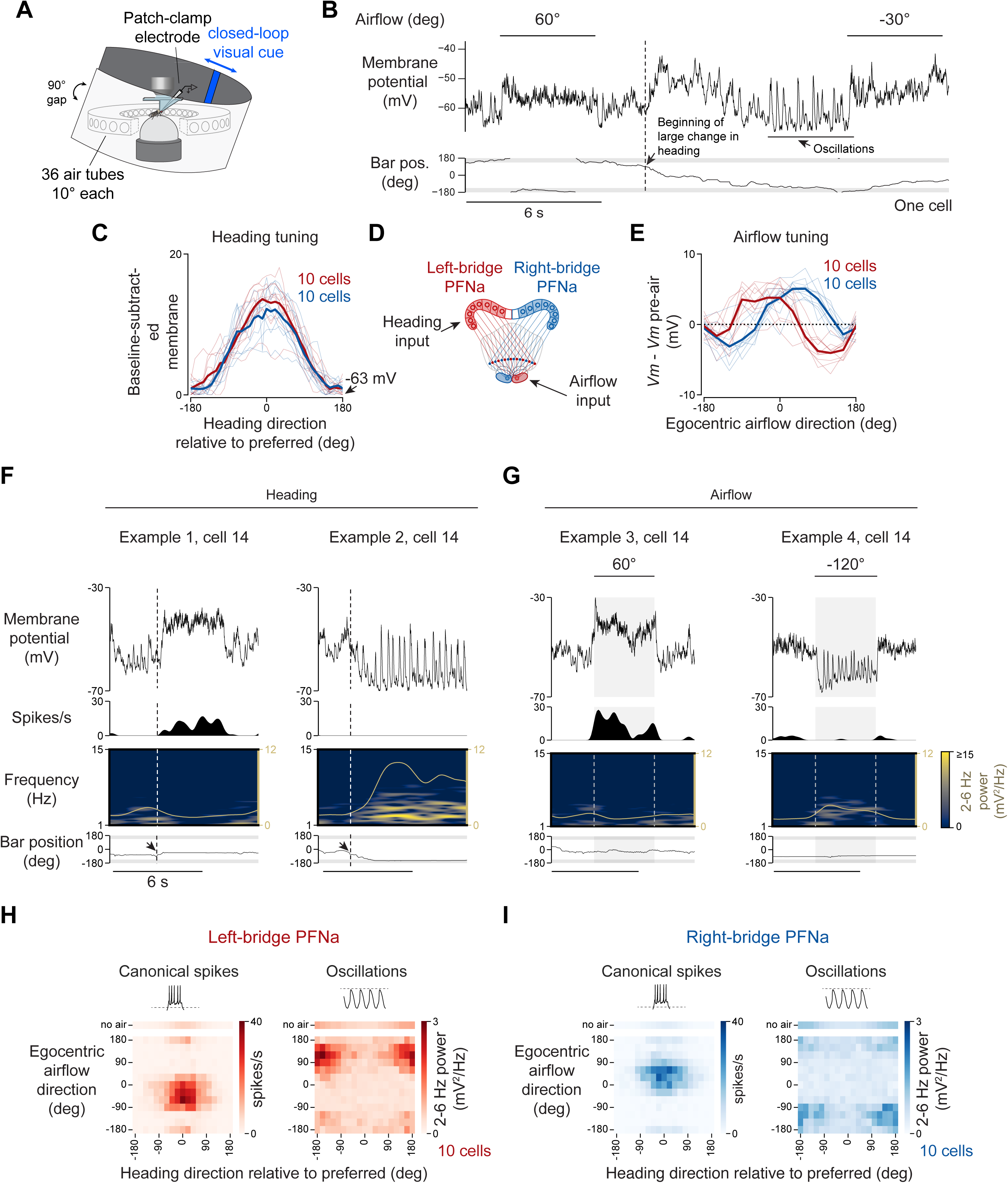
PFNa neurons fire canonical spikes when depolarized and express oscillations when hyperpolarized, with the oscillations signaling airflow stimuli from the egocentric rear. **(A)** Patch-clamp of PFNa neurons in a fly walking with a constant heading cue and intermittent air puffs. **(B)** Example membrane potential (*Vm*) of a right-bridge PFNa neuron as the fly changes heading and senses airflow. (C) *Vm* heading tuning curves, aligned to each cell’s preferred direction. The mean raw *Vm* at baseline was –63 mV; the minimum *Vm* of the curve was subtracted to normalize. **(D)** Inputs to the PFNa neurons. **(E)** Airflow direction tuning curves. **(F)** Example responses from the neuron in B when the fly changed heading (dotted lines). **(G)** Same as in F, but for airflow epochs. **(H and I)** Conjunctive tuning of left-bridge or right-bridge PFNa-cell activity to the direction of airflow and heading. The top row in each heatmap shows the spike rate (or oscillation power) without airflow.

Conjunctive tuning to allocentric heading and the egocentric airflow direction is expected from a set of PFN neurons that aim to transform the airflow direction experienced by a fly from egocentric to allocentric coordinates. However, these results were initially confusing because they revealed PFNa cells to have (spike removed) *Vm* tuning curves to airflow with only a single peak, at approximately ±45° (**Figure 3E**), whereas the same cells showed double-peaked tuning curves to the same stimuli when measuring calcium in the noduli (**Figure 2G**).

In the course of performing these experiments, we noticed that when PFNa neurons were strongly hyperpolarized, their membrane potential showed large oscillations with a frequency of ∼2-6 Hz^21^ (**Figure 3B, S4A and S4B**). Using power in the 2-6 Hz band as a quantitative measure of the oscillation strength, we noted robust *Vm* oscillations whenever the membrane potential of PFNa cells was sufficiently hyperpolarized, independently of whether the hyperpolarization was caused by the fly changing its heading (**Figure 3F**, right) or by an air puff (**Figure 3G**, right). Similarly, PFNa cells expressed canonical sodium spikes when their *Vm* was sufficiently depolarized, regardless of the cause of the depolarization (**Figures 3F and 3G**, left).

To depict how PFNa neurons respond to combinations of headings and airflow directions, we plotted two-dimensional tuning curves (i.e. heatmaps) of canonical spikes (**Figure 3H**) and oscillation strength (**Figure 3I**) as a function of these two variables. We observed the strongest oscillations in response to stimuli that induce the largest membrane hyperpolarization, that is, anti-preferred heading angles and ±120° airflow angles (**Figures 3H and 3I**, right panels). The airflow directions that elicited *Vm* oscillations matched those that yielded phase inversions of the PFNa sinusoids in the protocerebral bridge in relation to the EPG phase (**Figures 2F, 3H and 3I**). Conversely, the airflow directions that elicited the strongest canonical spikes in PFNa cells were those that yielded phase alignments between PFNa and EPG bumps in the bridge (**Figures 2E, 3H and 3I**). A switch in signaling from depolarization-driven canonical spikes to hyperpolarization-driven oscillations is thus poised to underlie vector inversions at the level of calcium signaling. These findings also explain how the LNOa inputs to PFNa cells, which express only a single calcium peak across airflow directions, can drive a two-peaked calcium response in PFNa cells. One calcium peak, matching the regime of low LNOa activity, was likely due to PFNa cells being depolarized and firing canonical sodium spikes; this sign makes sense since LNOa cells are expected to be inhibitory on the PFNa cells^21^ (**Figure S2F**). The other peak, matching the regime of high LNOa activity, was likely due to PFNa cells being sufficiently hyperpolarized to express oscillations, which could also lead to elevated calcium.

### Hyperpolarization-elicited oscillations in PFNa cells are mediated by Ca-**α**1T calcium channels

To test whether the second calcium peak in the PFNa noduli tuning curves stemmed from *Vm* oscillations, we wished to better understand the mechanism of oscillatory *Vm* dynamics. We reasoned that if artificially hyperpolarizing a single PFNa cell could reliably trigger oscillations, this would suggest a role for intrinsic membrane conductances in the phenomenon. Consistent with that notion, we were able to routinely induce oscillations in PFNa neurons by injecting hyperpolarizing current into these cells (**Figures S4C, S4D, and S4G**).

The oscillations in the *Vm* of PFNa neurons reminded us of the burst firing mode of mammalian thalamocortical neurons, where low-voltage activated, T-type calcium channels are key to the production of rhythmic spiking^13,14^ (**Figure 4A**). Specifically, hyperpolarization of thalamic neurons relieves T-type channels from inactivation, enabling them to produce regenerative calcium spikes^13,14^. The *Drosophila* genome encodes a single T-type calcium channel, Ca-α1T, and an RNA sequencing dataset^28^ revealed that PFNa cells express the *Ca-α1T* transcript at a 35-fold higher level than the median of all cell types analyzed in that study (**Figure 4B**). In addition, we observed strong immunohistochemical signal from a GFP-tagged knock-in allele of Ca-α1T, *GFP::Ca-α1T*^29^, in the third layer of the noduli and the ventral layers of the fan-shaped body, which are regions innervated by PFNa neurites (**Figure 4C**). These observations suggested that a Ca-α1T-mediated conductance may endow PFNa neurons with the ability to oscillate when hyperpolarized.

**Figure 4.**
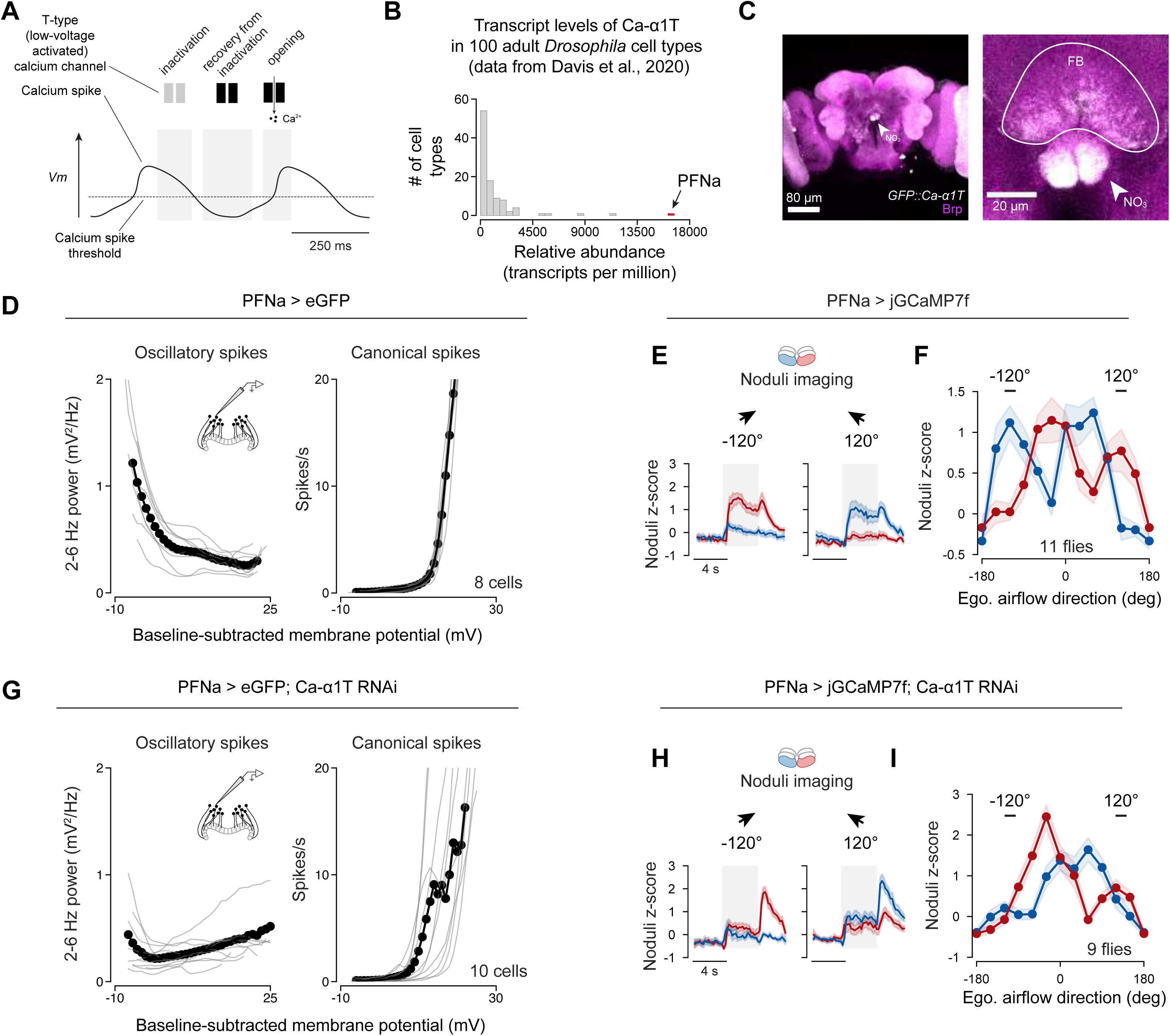
The T-type calcium channel Ca-a1T mediates oscillations in PFNa neurons and thus the ability of these cells to signal rear airflow stimuli. **(A)** How T-type calcium channels contribute to calcium spikes. **(B)** Relative abundance of *Ca-a1T* transcripts in a published dataset^28^. **(C)** Expression pattern of GFP::Ca-a1T^29^. Image is representative of 3 brains stained. **(D)** 2-6 Hz power (left) and canonical spike rate (right) as a function of *Vm* in control PFNa neurons. Data from left- and right-bridge neurons was pooled. Dots show the mean. **(E)** jGCaMP7f responses to airflow from ±120°, imaged in the noduli of control flies. The peak observed at airflow offset is consistent with post-inhibitory rebound. Mean and SEM shown. **(F)** Airflow tuning curves of jGCaMP7f responses at the noduli. Points show the average value between 2 and 4 seconds after airflow onset. Mean and SEM shown. **(G and I)** Same as D, F, but in PFNa cells with *Ca-a1T* knockdown.

We knocked down the transcript levels of Ca-α1T in PFNa neurons using RNAi^30^ and recorded PFNa *Vm* while presenting open-loop air puffs to flies navigating a virtual environment. Unlike in control flies, PFNa neurons with *Ca-α1T*-knockdown rarely expressed any *Vm* oscillations to airflow stimuli arriving from behind (**Figures 4D, 4G, S3G, S3H, S3I, and S3J**) or when hyperpolarized with current injection (**Figures S4E, S4F, S4H, and S4I**). PFNa neurons with *Ca-α1T*-knockdown also showed diminished secondary calcium peaks to airflow stimuli arriving from behind when imaging the noduli (**Figures 4E, 4F, 4H, and 4I**). In a second approach for abrogating Ca-α1T function, we patch-clamped PFNa neurons in Ca-α1T-mutant flies (Ca-α1T^del/Δ135^) flies and did not observe clear oscillations in response to hyperpolarization (**Figure S4K**). Together, these results demonstrate a critical role for T-type channels in generating *Vm* oscillations within PFNa neurons. We thus refer to these oscillations as *calcium spikes* hereafter; a characterization of these spikes is shown in **Figures S5A-S5I**. These results also implicate the *Vm* oscillations in generating the calcium signals associated with air puffs from behind, which underlie the PFNa system’s ability to invert vectors.

### A conceptual model for how PFNa neurons can invert their encoded vector

These findings led us to the following conceptual model for how a population of PFNa neurons can encode a vector whose direction is invertible. Each population of PFNa cells––one in the left bridge and the other in the right bridge––expresses a sinusoidally shaped *Vm* signal across eight glomeruli on its side of the protocerebral bridge (**Figures 5A and 5B**). A threshold exists in the system and calcium rises when the *Vm* of a PFNa cell deviates from this threshold in either direction. Calcium influx occurs with sodium spikes because the membrane is depolarized enough to presumably activate high-voltage activated (HVA) calcium channels. With calcium spikes, calcium influx occurs via the T-type channels directly, or, potentially also indirectly via the activation of HVA calcium channels with sodium spikes that sometimes ride on top of the calcium spikes. In either case, a calcium bump that encodes a vector is induced across the neuronal population (**Figure 5C**, left).

**Figure 5.**
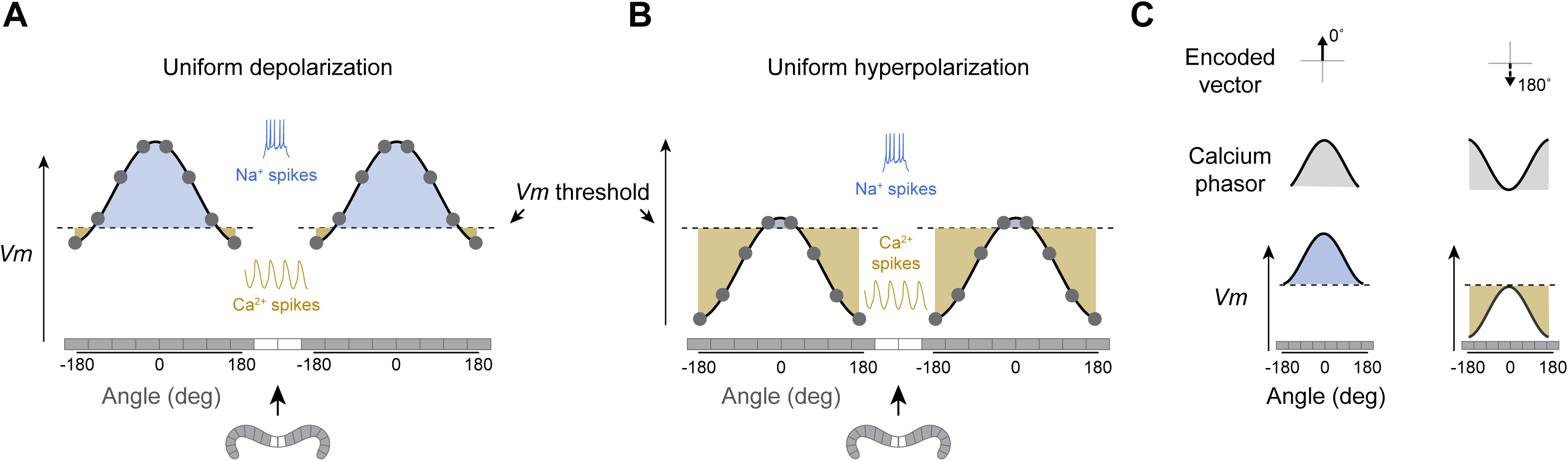
A qualitative model for vector computation with invertible vectors in PFNa neurons. **(A and B)** Sinusoidal population activity of a simplified set of 16 neurons tiling the protocerebral bridge. Cells more depolarized than a threshold fire sodium spikes, more hyperpolarized cells fire calcium spikes instead. The shaded area is the total sodium or calcium spike intensity across the population. **(C)** Relationship between Vm, population calcium signals, and encoded vectors. The higher above threshold that a neuron’s Vm is, the stronger it will fire sodium spikes and the more calcium will enter the cell. The expected population calcium signal and its encoded vector would point at 0°. With uniform hyperpolarization, the sinusoidal population calcium signal and its encoded vector point at 180°.

LNOa input to the PFNa population in the nodulus can uniformly depolarize (**Figure 5A**) or hyperpolarize (**Figure 5B**) the PFNa population as a function of the air puff direction. When the PFNa population is uniformly depolarized, the system expresses a sodium-spike induced calcium bump that is aligned with the EPG bump (set to be 0° in **Figure 5C**, left). When the PFNa population is uniformly hyperpolarized, cells 180° away from the EPG peak are maximally hyperpolarized and the system generates a T-type spike induced calcium bump whose peak is 180° offset from the EPG bump, thus implementing vector inversion (**Figures 5B and 5C**, right). In this way, an LNOa input signal with a single tuning peak can induce two peaks in the calcium tuning curves of PFNa cells––one peak due to sodium spikes and the other due to calcium spikes––while also inverting the vector encoded by the PFNa population across the two spike modes.

A key remaining question was whether postsynaptic neurons sum the phase-aligned and phase-inverted PFNa vectors to generate a 360° airflow-direction signal. Partly because T-type calcium currents are thought to have little direct impact on fast synaptic release^31^, we actually considered this possibility relatively unlikely. We reasoned instead that downstream neurons would primarily respond to the sodium spike–encoded vectors, with the calcium spike–encoded vectors serving a slower, modulatory role, perhaps evident only under specific conditions. To probe this matter, we examined how FC3 neurons—fan-shaped body cells postsynaptic to PFNa cells—read out the PFNa vectors, discovering that they do so with some flexibility.

### FC3 cells read out the EPG-phase-aligned vectors during air puffs

The sum of the left and right PFNa sinusoids should be another sinusoid that encodes a vector pointing in the allocentric direction of air puffs (**Figures 1B and 1G**). We tested whether FC3 neurons, which are monosynaptically downstream of PFNa cells^17^ (**Figure 6A** and **Figure S6A**), perform such a vector-sum operation on their PFNa inputs.

**Figure 6.**
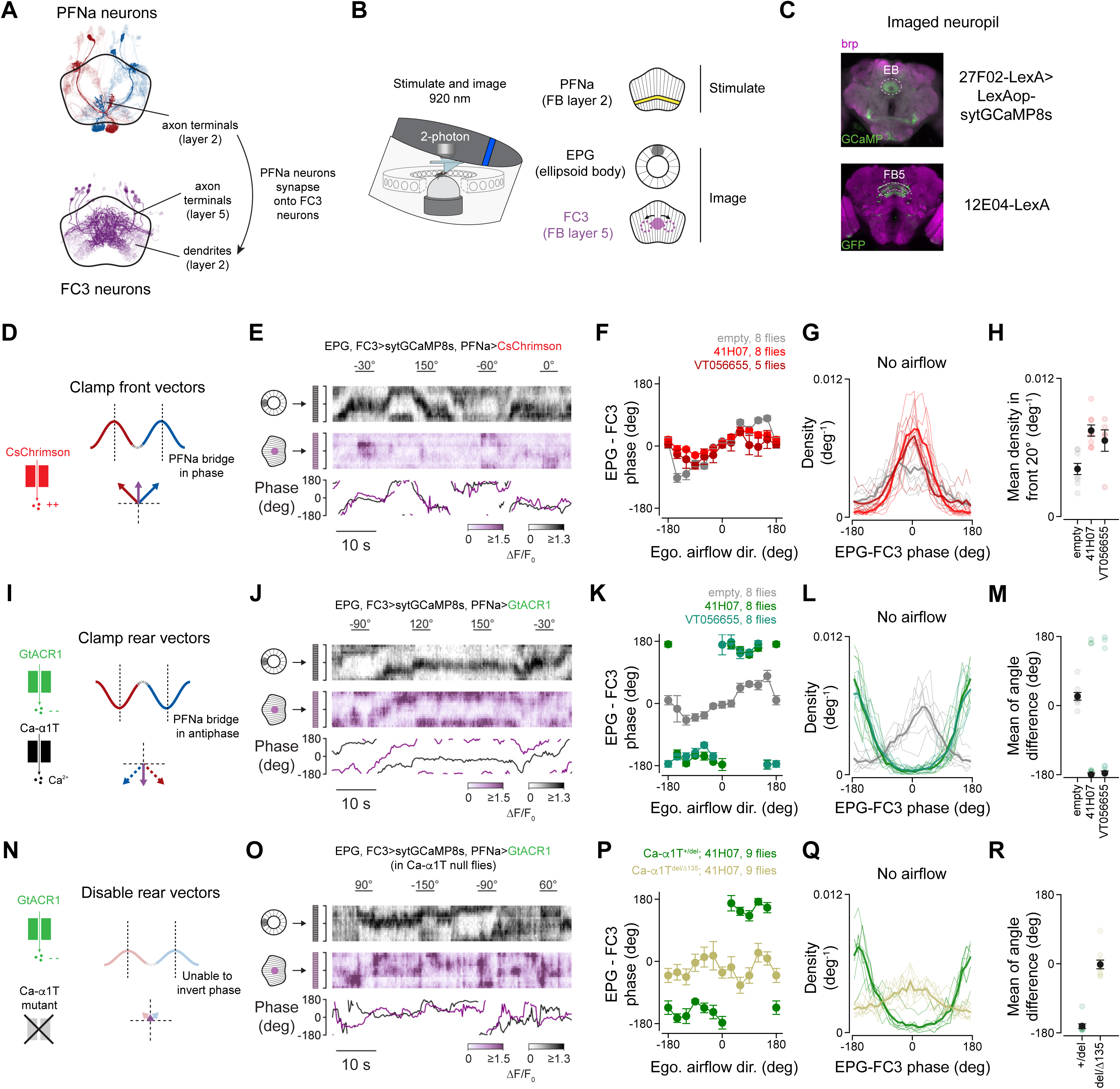
FC3 neurons can functionally sum two sodium-spike or two calcium-spike encoded vectors in the PFNa populations. **(A)** EM rendering of the PFNa and FC3 neurons. The dark-colored PFNa neurons provide input to the dark-colored FC3 neurons. **(B)** Simultaneous imaging of the EPG and FC3 neurons while stimulating the PFNa neurons. **(C)** Expression pattern of the driver lines used for imaging. **(D)** Expected PFNa activity in the bridge (red and blue) upon depolarization, relative to the EPG phase (dotted lines). **(E)** Example EPG and FC3 activity (heatmaps) and bump phases (bottom) in the context of PFNa optogenetic depolarization. **(F)** Mean EPG – FC3 phase difference in response to airflow stimuli during optogenetic depolarization of PFNa neurons. Mean and SEM shown. **(G)** Probability distributions of the EPG-FC3 phase difference without airflow and during optogenetic depolarization of PFNa neurons. **(H)** Mean density in the frontal 20° of the EPG-FC3 phase difference. Mean and SEM shown. **(I, J, K, L)** Same as in D, E, F, G, but for optogenetic hyperpolarization via GtACR1. **(M)** Circular difference of the average density of EPG-FC3 phase difference curves in K. Mean and SEM shown. **(N, O, P, Q, R)** Same as in H, J, K, L, M,, but in Ca-α1T^del/Δ135^ null mutants. Heterozygous flies (Ca-α1T^+/del^) are used as controls.

We imaged calcium in FC3 and EPG neurons (**Figure S6B**) while head-fixed flies navigated a virtual environment and received air puffs from twelve directions. Akin to EPG cells in the ellipsoid body, FC3 cells express a single bump of calcium activity whose phase in the fan-shaped body changes over time (**Figure S6B**). Without airflow, the phase of the FC3 bump was generally aligned to the EPG phase, though with frequent and sometimes quite large deviations from the EPG phase (**Figure S6B**, purple and black curves). During air-puff stimuli, the FC3 bump amplitude increased and its phase systematically realigned in reference to the EPG phase (**Figure S6C**) as a function of the egocentric angle of the airflow for puffs delivered within ∼±55° of the midline. These data are consistent with the PFNa front vectors––encoded by sodium spikes––being able to reposition the FC3 calcium bump in the fan-shaped body. For air puffs delivered more peripherally than ∼±55° from the midline (i.e., airflow from the sides and rear), the FC3 phase did not systematically deviate beyond ∼±55° (**Figures S6D and S6E**). With air puffs directly from the back, we did not observe a consistent difference between the EPG and FC3 phases (**Figures S6D and S6E**). These data suggest that the inverted, rear-facing vectors––encoded by PFNa calcium spikes––have a much weaker influence on the position of the FC3 bump.

To quantitatively address how much, if any, the calcium spikes contributed to the FC3 phase signal in these experiments, we used the sodium spike-rate and oscillation-power data from the electrophysiological recordings of PFNa cells (**Figures 3H and 3I**) to predict the FC3 phase. In brief, we summed a normalized version of the sodium-spike and oscillation data, shifting the heading angle appropriately for left and right PFNa neurons based on how PFNa neurons project from the bridge to the fan-shaped body; the shift angles were either ±39°, ±45°, or ±55° so we could test the full range of potential deviations that this system can accommodate (**Methods**). We scaled the oscillation-power signal by a free parameter and determined the phase of the summed output signal as a function of the airflow direction. For all PFNa shift angles tested, a ∼10% contribution for the oscillation-power signal was needed to attain the best fit, which was always considerably better than the fit obtained with a 0% contribution from the calcium oscillations (**Figures S6E-S6G**). These data-driven fits suggest that the calcium-spike-based rear vectors contribute to positioning the FC3 phase in the fan-shaped body, albeit with a considerably smaller weight than the sodium-spike-based front vectors. The fact that ∼4% of calcium spikes had sodium spikes riding on top them in the context of these experiments (**Figure S5**)––and such sodium spikes are expected to induce fast synaptic transmission––provides a viable mechanism for how a subset of calcium spikes might have contributed to a postsynaptic FC3 response. We interpret these data to mean that the PFNa calcium spikes sometimes induce an immediate postsynaptic response, perhaps when they have sodium spikes riding on top (**Figure S5C**), but that this might not be their primary function. We speculate that the calcium spikes also implement a process that is separable from the immediate release of classic synaptic vesicles, perhaps modulating synaptic weights on longer timescales –– an idea we return to in the Discussion.

### FC3 cells read out the EPG-phase-aligned and EPG-phase-inverted PFNa vectors during optogenetic stimulation of PFNa cells

Our data-driven fits suggested a weak association of calcium spikes with fast synaptic transmission (**Figure S6G**). However, we occasionally observed large deviations of the FC3 phase relative to the EPG phase, well beyond what summing the sodium-spike-based PFNa vectors could support (e.g., **Figure S6B**, arrows). These observations suggested that there might exist conditions in which the calcium-spike-based PFNa sinusoids elicit strong and reliable phase shifts in FC3 neurons just like the sodium-spike sinusoids. We uncovered such a situation using optogenetics.

According to our model, uniform depolarization of both PFNa populations should extend the two EPG-phase aligned vectors. If the activity of FC3 neurons reflects the sum of these two vectors, the phase of the FC3 bump should align with the EPG phase during such PFNa depolarization. To test this hypothesis, we optogenetically depolarized PFNa cells using CsChrimson^32^ while simultaneously imaging the FC3 and EPG calcium bumps (**Figures 6A-6D**). We found that with optogenetic depolarization of PFNa cells, the FC3 and EPG phases became more aligned (**Figures 6D-6H**), consistent with our model. This effect was clear both during airflow (**Figure 6F**)––where we observed muted deviations of the FC3 phase from the EPG phase for air puffs from the sides and rear––and during moments when no airflow was presented (**Figures 6G and 6H**).

We also performed the converse experiment, optogenetically hyperpolarizing PFNa cells with the chloride-selective channelrhodopsin GtACR1^33^ while simultaneously imaging the FC3 and EPG calcium bumps (**Figures 6I-6M**). Uniform hyperpolarization of both PFNa populations should induce calcium spikes in PFNa cells, thus extending the two EPG phase-inverted vectors (**Figure 6I**). If such a perturbation were to elicit synaptic transmission from PFNa to FC3 cells, the long, phase-inverted PFNa vectors should yield a downstream FC3 bump whose phase is 180° offset from the EPG phase. We found that with PFNa cells optogenetically hyperpolarized, the FC3 bump was consistently antiphase to the EPG bump (**Figures 6I-6M**), providing evidence for the ability of the calcium-spike-associated vectors to impact downstream physiology. This EPG-FC3 antiphase effect was clear both during air puffs (**Figure 6K**) and at moments when no air puffs were presented (**Figures 6L and 6M**).

Eliminating the ability of neurons to express T-type calcium spikes should eliminate phase-inversion of the FC3 bump relative to the EPG bump with PFNa hyperpolarization (**Figure 6N**). We repeated the hyperpolarization experiment in in Ca-α1T-null flies (Ca-α1T^del/Δ135^), which lack functional T-type calcium channels^29,34^ and hyperpolarization induced T-type spikes in PFNa neurons (**Figure S4K**). Consistent with our model, the FC3 bump no longer appeared phase inverted relative to the EPG bump in these experiments (**Figures 6O-6R**). These results demonstrate that PFNa calcium spikes can induce strong postsynaptic FC3 activity and that this FC3 activity can reflect a sum of the PFNa input vectors when the two input vectors are phase-inverted to the EPG bump and of equal length. We verified that optogenetic hyperpolarization of PFNa neurons induces T-type calcium spikes in non-mutant flies (**Figure S4J**). In the Discussion, we consider why the calcium spikes might have induced robust transmitter release in the optogenetic but not the air-puff experiments. Taken together, our optogenetic results demonstrate that both EPG phase-aligned and EPG phase-inverted vectors in PFNa cells can induce vector-addition-like signaling in downstream cells.

### A quantitative model for computing allocentric directions using invertible vectors

Inspired by the conceptual model and experimental results, we developed a mathematical model of the PFNa system (**Figure 7A**). In the model, the PFNa bumps or phasors in the protocerebral bridge can exist both in phase and ∼180° out of phase with the EPG heading angle, thus instantiating invertible vectors. PFNa cells send axons from the bridge to the fan-shaped body with an offset that rotates the left- and right-bridge phasors by ±45°, giving rise to a pair of orthogonal vectors in the fan-shaped body that form a basis for the allocentric direction of airflow (**Figure 7A**). The allocentric airflow direction can be computed by summing the left and right PFNa outputs. Air puffs from the front lengthen the ±45° vectors represented by these outputs. Air puffs from the rear invert the vectors such that they shift to pointing at ±135° (**Figure 7A**), which allows the system to represent directions that have negative projections along the ±45° axes. We present a detailed analysis of these vector angles in the **Methods** and **Figure S7A-S7G**.

**Figure 7.**
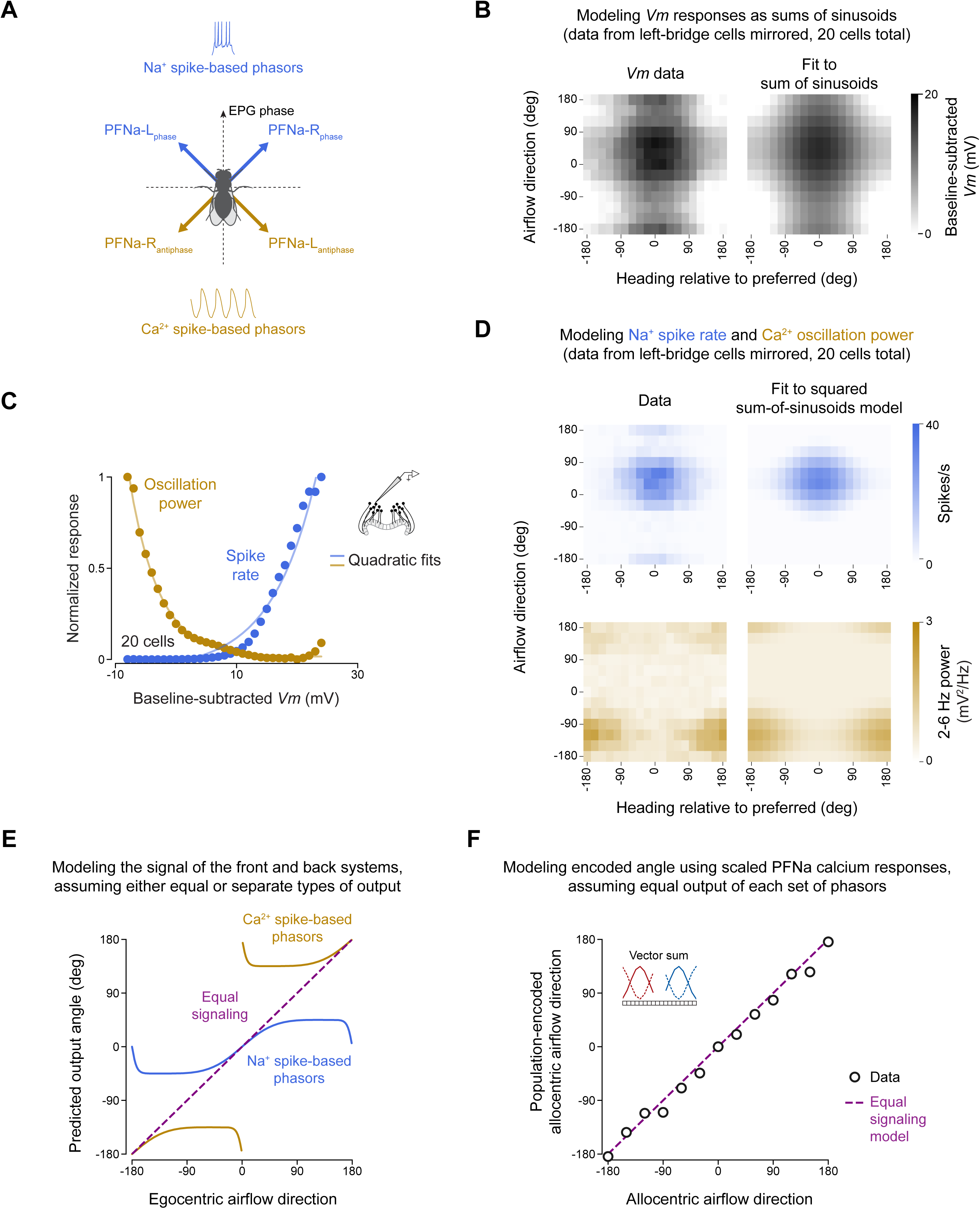
A quantitative model for vector computation with invertible vectors in PFNa neurons. **(A)** The PFNa vector system. Angles in the fly’s egocentric front are signaled by the sodium-spike-based PFNa vectors. Angles in the fly’s egocentric rear are signaled by the calcium-spike-based vectors. Normalized Vm responses of the PFNa neurons (left) and the fit to a sum of two sinusoids representing heading and airflow (right). Data is aligned to the cell’s preferred heading; data for the left-bridge cells was mirrored along the airflow axis to allow pooling. **(C)** Sodium spike and oscillation power responses of the PFNa neurons as a function of normalized membrane potential. Each curve has been normalized to values between 0 and 1 to highlight their relationship despite the difference of their units (spikes/s and 2-6 Hz power, respectively). Mean (dots) and quadratic fits are shown. **(D)** Sodium-spike and oscillation power responses of the PFNa neurons (left) and the prediction of the quadratic model (right). Data was heading-nulled and mirrored as in B. **(E)** Output of the quadratic model for the sodium-spike-based vectors only (blue), calcium-spike-based vectors only (gold), or a scenario where both types of vectors can add linearly (purple). **(F)** Predicted output direction of the PFNa neurons using the calcium responses in S2A (circles) and the quadratic model (purple). The phase-aligned and phase-inverted PFNa signals have different calcium sources and thus jGCaMP7f responses of different amplitude; we thus scaled the phase-inverted calcium responses by a uniform factor of 0.2 in this panel.

We modeled the PFNa membrane potential as a weighted sum of two terms: a cosine function of the airflow direction angle and a cosine function of heading. The quality of the resulting fits (**Figure 7B**, 92% and 94% variance explained) suggests that airflow and heading inputs combine additively. To fit the spiking responses of the PFNa cells (**Figures 3H and 3I**), we noted that both the sodium-spike rate and the calcium-oscillation strength show a quadratic dependence on *Vm* (**Figures 7C, S3C and S3F**). Using the sum-of-sinusoids model for *Vm*, followed by a squaring operation, we fit both the sodium-spike rate and the calcium oscillation strength across all airflow and heading angles (**Figure 7D** and **Methods**). The model explained 89% of the variance for the sodium spikes and 77% for the calcium spike oscillation strength (averaged across left and right PFNa neurons), suggesting that it captures much of the underlying physiology.

The experiment in **Figure S6** revealed that, in the context of open-loop airflow, the calcium-spike encoded vectors have a relatively small impact on downstream FC3 neurons. However, the experiment in **Figure 6** (panels **I-M**) revealed that calcium spikes, when induced optogenetically, have a strong impact on downstream FC3 neurons. Our formal model allows us to capture any arbitrary level of sodium- and calcium-spike based impact on FC3 cells (**Figure 7E**). When calcium spikes are dominant over sodium spikes, the model’s two-vector sum effectively tracks only angles in the rear (**Figure 7E, gold**) and when sodium spikes are dominant over calcium spikes the system effectively tracks only angles in the front (**Figure 7E, blue**). When the level of interaction is equal between the two signaling modes, the sum of the two vectors can comprehensively track all allocentric angles (**Figure 7E, purple**).

Our model allows us to understand why this is the case. Both the sodium spike rate and oscillation power can be described by rectified quadratic functions of the membrane potential with a similar threshold (**Figures 5 and 7C**). If these two modes contributed equally, the two rectified quadratic functions could be summed into a single unrectified squaring function, providing an ingenious way for a neural system to perform a full (unrectified) squaring operation. We have shown that the PFNa membrane potential is a sum of sinusoidal functions of heading and airflow direction (**Figure 7B**) and squaring this sum produces a phasor that precisely represents the allocentric airflow direction (**Methods**). The fact that we can decode an allocentric direction by assuming equal output from the sodium-spike and calcium-spike systems highlights that the two types of phasors share a reference frame. This shared reference frame means that even in contexts where the calcium-spike signal is not transmitted to downstream neurons, it could instead seed a biochemical process within the PFNa neurites whose rate or magnitude relies on vector operations –– an intriguing idea to be explored in future work.

## Discussion

### Function of the PFNa cells

PFNa neurons combine allocentric heading inputs from the protocerebral bridge with egocentric airflow-direction inputs from the noduli. Two PFNa populations, originating in the left and right bridge, generate sinusoidally modulated calcium signals that function as invertible vectors. By summing these two invertible vectors the allocentric direction of airflow (or, potentially, other directional stimuli that activate PFNa cells) can be calculated (**Figure 7F**). In situations where flies are standing still, or walking slowly, the direction of airflow sensed by the body directly reflects the direction of wind in the external world. The direction of wind is of broad importance to navigating insects^9,35–42^, and thus this circuit has the potential to generate a signal^19,43^ that can guide many homing and food-orienting behaviors that rely on wind assessments^21^. Our results tentatively speak against the idea that PFNa neurons––unaided by other PFN cells––routinely build a pure, 360°-spanning allocentric wind signal in postsynaptic neurons. However, it is possible that the PFNa signals contribute to airflow orientation in ways other than building a postsynaptic calcium bump that is strictly analogous to the hΔB traveling direction signal.

### Synaptic transmission and T-type calcium spikes

Classic synaptic release relies on calcium entry through high-voltage-activated calcium channels, which are found very close to the synaptic-vesicle fusion machinery^44,45^. Because low-voltage activated calcium channels, like T-type channels, do not typically reside immediately adjacent to active zones, calcium entry through these channels is not commonly thought to drive vesicle fusion. The air-puff responses of FC3 neurons, which are monosynaptic recipients of PFNa input, generally support this dichotomous view of calcium signaling at synapses. That is, the calcium bump of the FC3 neurons is consistently repositioned by the sodium-spike-based vectors, but only rarely by the calcium-spike-based vectors. In contrast to airflow, optogenetically hyperpolarizing the PFNa neurons was able to induce a an FC3 bump 180° offset from the EPG bump. What could have been the difference between the airflow and optogenetic experiments? One possibility is that a specific modulator state is needed to yield synaptic transmission of the calcium-spike encoded vectors, perhaps by increasing the likelihood that sodium spikes ride on top of the calcium spikes (**Figures S5A-S5J**). Optogenetic stimulation may have been sufficiently strong so as to bypass the need for such a modulatory input. (Because the amount of hyperpolarization that chloride channels, like GtACR1, can evoke is limited, our optogenetic stimulation is likely to have induced a strong, but still physiologically relevant state).

Our results to date are agnostic as to whether the central complex can combine one sodium-spike (phase-aligned) phasor (say, in the left bridge) with one calcium-spike (phase-inverted) phasor (say, in the right bridge) to perform a vector sum. Our quantitative model argues that this sort of interaction has the potential to generate an accurate allocentric angle as its output––with appropriate weighing of the two signals––but our air-puff experiments suggest that these two modes of signaling may not directly sum. Indeed, the wide (∼200 ms) calcium spikes elicited by T-type calcium channels seem better suited for promoting peptide release from dense core vesicles^46–48^, and the release of neuropeptides into the fan-shaped body might not be immediately apparent at the level of postsynaptic calcium. Neuropeptides may elicit downstream molecular processes that are sinusoidally modulated in their intensity across the left/right extent of the fan-shaped body, representing a vector memory in the system that can alter navigation-related computations at a later timepoint.

### Vector integration and T-type calcium spikes

Rhythmic T-type calcium spikes in PFNa neurons are also well-poised to serve intracellular signaling roles, rather than, or in addition to, synaptic transmission-related roles. For example, calcium-calmodulin dependent protein kinase II (CaMKII) is particularly sensitive to calcium oscillations at the 2-6 Hz frequency range^49^, which matches the calcium spike rate of PFNa neurons. Because the calcium spike amplitude (i.e., the power of the 2-6 Hz *Vm* oscillation) varies sinusoidally across the left/ right extent of the fan-shaped body, an integral of this process could represent a vector that grows in amplitude with each T-type spike across the population. Such an integrated vector in the PFN axons^16^ could indicate for how long airflow has arrived at the fly’s rear, for example, which might be useful for driving orienting behaviors. Calcium potentials could also mark synapses as being eligible for plasticity^50^, or serve to adjust the rate of biochemical integration of sodium spikes, creating a sinusoidally modulated vector-trace signal across the fan-shaped body in this manner as well.

### Neural computation and T-type calcium spikes

Calcium signals in neurons are commonly used as a proxy for the sodium spike rate and presumed synaptic output. Our study adds nuance to this assumption. High calcium levels in PFNa neurons can indicate either a high sodium-spike rate or a high calcium-spike rate, yet only the former reliably drives synaptic output. Whereas calcium spikes are mainly relevant for cells with T-type calcium conductances, expression of this channel family is widespread across animals^51^. Beyond flies, broad calcium spikes have been observed in the giant motor axons of the jellyfish *Aglantha*^52^ and the neurons of the mammalian inferior olive^53^. Delta rhythms in thalamocortical networks, which rely on T-type calcium channels^54,55^, are a hallmark of sleep^14^. Hippocampal and cortical networks express oscillatory dynamics in the delta and theta range during navigational tasks, and the functions of these oscillations are still being studied^56–58^. Our work shows that T-type calcium channels can produce quantitatively precise vector signals in *Drosophila*. Similarly explicit, real-time computational functions for calcium spikes likely await discovery in other systems.

### Limitations of the study

Our study revealed that T-type spikes allow PFN cell populations to invert their vector signals, but how these signals contribute to natural behavior remains unclear. How FC1, hΔC, hΔJ, and FR1 cells––the other major downstream partners of PFNa cells––read out the PFNa vector signals remains unknown. We observed sodium spikes that were not predicted by our model in response to air puffs from 180°; studying the PFNa system in the context of a closed-loop wind task could resolve whether those spikes represent anomalous responses to open-loop air puffs or instead are a critical component of how the system naturally works.

## Resource availability

### Lead contact

Requests for further information and resources should be directed to and will be fulfilled by the lead contact, Gaby Maimon.

### Materials availability

All unique/stable reagents generated in this study are available from the lead contact without restriction.

### Data and code availability

- Datasets associated with this paper are available at https://doi.org/10.5281/zenodo.17555686 and are publicly available as of the date of publication.
- Code for data analysis and modeling, as well as experiment scripts and 3D models of the airflow device and associated custom control box, are available at https://github.com/MaimonLab/cx-vector-inversion is publicly available as of the date of publication.
- Any additional information required to reanalyze the data reported in this paper is available from the lead contact upon request.

## Acknowledgements

We thank Jim Petrillo at the Precision Instrumentation Technologies facility at Rockefeller University for extensive assistance with the design and fabrication of the static airflow device, as well as Andy Siliciano, Chad Morton, and Vanessa Ruta for sharing designs for an airflow delivery device^59^ upon which ours is based; Jazz Weisman for help with initial prototypes of the static airflow device; Cheng Lyu and Stephen Thornquist for the LexAop-syt-jGCaMP8s stock, Stephen Thornquist for the UAS-GFlamp1 stock, and Yulong Li for the UAS-GFlamp1 plasmid; Peter Mussells Pires for scripts for segmenting ROIs, advice on rendering neuron skeletons from the hemibrain connectome, and comments on the manuscript; Audrey Francis and Victoria Parache for assistance with immunostaining; Abigail Janke for advice on driver line selection; and the entire Maimon laboratory for helpful discussions. Research in this manuscript was supported by a Kavli postdoctoral fellowship to S.S. and a grant from the National Institute of Neurological Disorders and Stroke (R35NS132252) to G.M. L.F.A was supported by The Gatsby Charitable Foundation, The Kavli Foundation, and the Simons Collaboration for the Global Brain. G.M. is a Howard Hughes Medical Institute (HHMI) Investigator.

## Author contributions

I.G.I. and G.M. conceived of the project. I.G.I. performed the imaging and optogenetic experiments and the associated analyses. S.S. performed the electrophysiological experiments and the associated analyses. M.H. performed the current injection experiments in Ca-α1T^del/Δ135^ flies. I.G.I, S.S., L.F.A. and G.M. jointly interpreted the data and decided on new experiments. L.F.A. developed and implemented the formal models. T.L.M. and I.G.I. developed the airflow device. I.G.I., L.F.A., and G.M. wrote the paper with input from S.S. and T.L.M.

## STAR Methods

### Experimental model and study participant details

#### Fly husbandry

Unless indicated otherwise, flies were reared in standard cornmeal-agar-molasses food in a 12h/12h light cycle incubator set to 25 °C. Progenies from crosses were transferred into fresh vials on the day of eclosion and housed in an incubator for 2-7 days before being affixed to a physiology platform for calcium imaging. For electrophysiological experiments, we used 4-7 day old flies. All animals used in this study were adult female *Drosophila melanogaster*.

Fly crosses for optogenetic experiments were raised in cornmeal-agar-molasses containing vials, wrapped in aluminum foil to minimize light exposure during development. On the day of eclosion, newly hatched flies were transferred into cornmeal-agar-molasses containing vials supplemented with 0.4 mM all-trans retinal (Sigma Aldrich). These vials were wrapped in aluminum foil for 2-5 days, until flies were affixed to a physiology platform for imaging experiments.

#### Fly stocks

Genotypes for each experimental cross are listed in **Table S1**; Stock sources are listed in the Key Resources Table.

We used existing Gal4 and LexA driver lines^64–68^ to target transgene expression to central-complex neurons. We used the 12E04-LexA driver line to target FC3 neurons, and we used the 27F02-LexA, 60D05-LexA, and 60D05-Gal4 lines to target EPG neurons. In addition to targeting EPG neurons, we found that the 60D05-LexA and 60D05-Gal4 lines also target unidentified cells that innervate layers 2 and 5 of the fan-shaped body, which have airflow responses (data not shown). Thus, whenever we used 60D05-Gal4 or 60D05-LexA to image EPG neurons in conjunction with either PFNa or FC3 neurons, we expressed the red calcium indicator jRGECO1a^69^ to avoid uncertainty as to the cellular origin of fluorescence signals in the fan-shaped body. In optogenetic experiments––where it was not possible for us to work with two different calcium indicators––we used 27F02-LexA to drive syt-jGCaMP8s (based on jGCaMP8s^70^ and described below) expression in EPG cells because this driver line does not target airflow-responsive fan-shaped body cells. For all other imaging experiments, we used jGCaMP7f^71^ instead.

We targeted PFNa neurons using four different driver lines: the split-Gal4 SS02255^72^, 30E10-Gal4, 41H07-Gal4, and VT056655-Gal4. From inspection of the publicly available multi-color Flp-out images by Janelia Research Campus^73^ as well as from our own immunohistochemistry data, we believe that PFNa neurons are the only PFN cells targeted in these four driver lines. We found that using 30E10-Gal4 to drive UAS-GtACR1 expression was lethal at the pupal stage, and thus we used 41H07-Gal4 and VT056655-Gal4 for the experiments in **Figure 6**. We targeted the LNOa neurons using split-Gal4 SS047432^72^.

We designed the syt-jGCaMP8s construct by linking the *Drosophila* synaptotagmin-1 coding sequences and jGCaMP8s (Addgene Plasmid #162380)^74^ using a GSGSGS linker, with the Syt1 sequence at the N terminus of jGCaMP8. We then placed this construct into pJFRC19-13xLexAop2 backbone (Addgene Plasmid #26224)^64^, replacing myrGFP with syt-jGCaMP8s. The plasmid was synthesized by GenScript and inserted in the VK00022 landing site (BDSC Stock #9740) using PhiC31-based integration, performed by BestGene.

We created the recombinant stock 27F02-LexA (attp40), LexAop-syt-GCaMP8s (VK00022) to image the activity of the heading system in the ellipsoid body in Figure 6. Whereas 27F02-LexA normally targets the EPG neurons alone within the central complex, and the expression pattern and physiology of this stock is consistent with EPG targeting, we found that this specific recombinant stock also displayed faint GCaMP expression in central complex regions not targeted by the EPG cell class, namely, the outermost two glomeruli of the protocerebral bridge and nodulus 1. These two regions are diagnostic of PEN neurons and thus 27F02-LexA (attp40), LexAop-syt-GCaMP8s (VK00022) is likely to weakly target PEN1 or PEN2 cells in addition to EPG neurons. The phase signals of the EPG, PEN1, and PEN2 neurons are generally aligned in the ellipsoid body (the relevant neuropil in Figure 6), and thus ––other than brief and small phase offsets between these bumps at moments when the fly turns^24,25^ ––we do not expect the potential contribution from PEN neurons to the measured EPG phase estimates in Figure 6 as a concern for the conclusions drawn.

10xUAS-GFlamp1 was created using an attB-site carrying plasmid gifted to us by Yulong Li’s research group^75^. We inserted the plasmid at the VK00005 integration site (BDSC #9725) with PhiC31-based integration, performed by BestGene.

### Method details

#### Fly mounting

We cold-anesthetized and mounted adult female flies to a custom stage, which allows for head-fixed behavior simultaneous with neural imaging as described previously^76^. In brief, we attached the dorsal tip of the head and the anterior tip of the thorax to a form-fitting hole in the stage using a blue-light-activated glue (Bondic). After being thus attached, the posterior edge of the head capsule can be dissected for physiological measurements from the brain. For calcium imaging experiments that did not require optogenetic activation of neurons, we allowed the flies to recover for ≥ 2 hours under low levels of ambient light after being mounted. For experiments with imaging and optogenetics (**Figure 6**), we allowed the flies to recover for ≥ 2 hours inside a dark cardboard box. For electrophysiology experiments, we allowed the flies to recover for 2-4 hours inside a dark cardboard box. We used slightly different head pitch angles, depending on the brain structure which we needed to access. The angle between the front vertical drop of the thorax and back of the fly’s head in experiments that involved co-imaging the ellipsoid body and the fan-shaped body was ∼60° (as in refs. ^11,77^). This same angle was closer to ∼45° for experiments in which we performed imaging of the protocerebral bridge and the noduli, or electrophysiology from PFNa somas.

#### Extracellular saline composition and delivery

For both imaging and electrophysiology experiments, we exposed the dorsal surface of the fly’s brain by cutting a rectangular window in the head capsule using a 30-gauge syringe (BD PrecisionGlide). We perfused the brain with an artificial extracellular saline solution^78^ bubbled with carbogen (95% CO_2_ / 5% O_2_). The composition of the saline solution, in mM, was 103 NaCl, 3 KCl, 5 N-Tris(hydroxymethyl) methyl-2-aminoethanesulfonic acid, 10 trehalose, 10 glucose, 2 sucrose, 26 NaHCO_3_, 1 NaH_2_PO_4_, 1.5 CaCl_2_, and 4 MgCl_2_. All chemicals were sourced from Sigma Aldrich. The solution’s osmolarity was measured to be ∼280 mOsm, and after carbogen bubbling, the solution’s pH was close to 7.3. The saline was delivered to the brain using a gravity-fed perfusion system. Using a Peltier device (SC-20, Warner Instruments) regulated by a closed-loop temperature controller (CL-100, Warner Instruments), we set the saline’s temperature, measured in the bath, to 22°C for calcium imaging experiments and 25°C for electrophysiology experiments.

#### Two-photon calcium imaging

Calcium imaging data were acquired using an Ultima IV two-photon microscope (Bruker) and a Chameleon Ultra II Ti:Sapphire tunable laser (Coherent). In experiments where GCaMP fluorescence was imaged alone, the laser wavelength was set to 925 nm. In experiments where we simultaneously imaged GCaMP and jRGECO1a (**Figures 2 and S2**), the Chameleon laser wavelength was set to 1000 nm and supplemental excitation of the jRGECO1a calcium sensor was provided by a coaxial second laser set to 1070 nm (Fidelity-2, Coherent). The power of the Chameleon laser, as measured at the back aperture of the objective, was 20-50 mW for all experiments. Emitted light was collected through a 40x/0.8 NA objective (LUMPLFLN 40XW, model 1-U2M587, Olympus), split by a 575 nm dichroic mirror (575dcxr, Chroma), and collected by a pair of GaAsP photomultiplier tubes (H7422P-40, Hamamatsu). The green channel, capturing the jGCaMP7f or jGCaMP8s emission signal, was filtered through a 490-560 nm bandpass filter (525/70m-2p, Chroma). In dual imaging experiments, the red channel (containing the jRGECO1a signal) was filtered through two stacked 590-650 nm bandpass filters (620/60m-2p, Chroma). The objective was mounted on a Piezo device (525800-400, Bruker), which allowed for rapid scanning along the z axis. We imaged exclusively in galvo-galvo mode and used the Piezo device to acquire imaging volumes consisting of 3 to 6 z-planes scanned at 128×128 pixel resolution. The volumetric scanning rates ranged from 3 Hz––for experiments with a large region of interest (ROI), such as when we co-imaged the fan-shaped body and ellipsoid body (e.g., **Figures 6 and S6**)–– to 8 Hz when we employed very small ROIs, like when imaging the noduli (e.g., **Figures 2 and S2**). The dwell time for each pixel ranged from 3.6 to 4.8 µs. These settings minimized fluorophore bleaching while still providing adequate fluorescence signals.

#### Optogenetic stimulation

We used the two-photon laser to perform simultaneous two-photon imaging and GtACR1^33^ or CsChrimson^32^ activation (**Figure 6**), similar to previous approaches^11,79,80^. In these experiments, the laser power was increased from 20 to 50 mW across the six imaging planes at increasing depth, following an exponential trajectory. The 20-mW plane covered the posterior end of the fan-shaped body, and the 50-mW plane covered the anterior end of the ellipsoid body. Such an exponential power increase had several advantages. First, by having the power in the fan-shaped body be relatively low, this limited photobleaching of jGCaMP8s in the fan-shaped body. Second, the low laser intensity in the fan-shaped body should help to lead to moderate, hopefully physiological, excitation levels of optogenetic reagents in PFNa terminals, rather than overactivation. Third, the high intensity in the ellipsoid body helped to increase the quality of the signal from this deep structure. We imaged volumes at ∼3.5 Hz in optogenetic experiments. Approximately three planes were dedicated to imaging the fan-shaped body and approximately three planes were dedicated to imaging the ellipsoid body. Because we were optogenetically activating the terminals of PFN neurons in the fan-shaped body, we thus activated PFN cells at ∼3.5 Hz and with a ∼50% duty cycle.

#### Electrophysiology

We cold-anesthetized flies and affixed them to a custom stage as described above and previously^76^. We opened a small cuticular window over the central complex using a 30-gauge syringe (BD) and removed the underlying fat and tracheal tissue, with fine forceps, to expose the brain. We illuminated the fly using an 850 nm LED (M850F2, Thorlabs) coupled to a 400 µm wide fiber optic cable (M28L01, Thorlabs) that was focused onto the fly with lenses (MAP10100100-A, Thorlabs). We visualized GFP-expressing cell bodies via standard epifluorescence, except that we used a custom GFP emission filter, which passed 510-560 nm and >800 nm (Chroma). This filter allowed us to visualize green and infrared while also cutting out the red fluorescence of Alexa-568, which we often included in our pipette for anatomical fills.

We pressure ejected 0.5% collagenase IV (Worthington) in extracellular saline from a pipette with a 4-6 µm tip positioned over the neural lamella and perineurial sheath, which weakened and ultimately breached these layers. We raised the bath temperature to ∼30°C during this desheathing process, which took < 5 min., to activate the collagenase. Following rupture of the perineurial sheath, we lowered the bath temperature to 21°C for ≥ 5 min. and increased the flow rate of the perfusion to ensure that the collagenase was fully washed out. We then performed additional manual clearing of tissue with the micropipette containing extracellular saline until individual somas of interests we exposed. We held the temperature of the bath between 24°C - 28°C for the remainder of the experiment.

We used borosilicate glass capillaries (BF150-86-7.5, Sutter Instruments), pulled on a P-1000 puller (Sutter Instruments) and fire polished with a microforge (MF2-LS2, Narishige) to produce pipettes with 6-12 MΩ resistance and ∼1 µm tip openings. Pipettes were filled with intracellular saline whose composition, in mM, was 140 potassium aspartate, 1 KCl, 10 HEPES, 1 EGTA, 0.5 Na_3_GTP, 4 MgATP, 13 biocytin hydrazide, and 0.02 Alexa Fluor 568 hydrazide (A10437, ThermoFisher Scientific). The pH of the intracellular saline solutions was adjusted to 7.3 with KOH and its osmolarity was adjusted to 265 mOsm with water.

We visualized somas for targeting with a patch pipette using a Kinetix sCMOS camera (Teledyne Photometrics) mounted on an upright epi-fluorescence microscope (Slicescope, Scientifica) with a 40×/0.80 NA water immersion objective (LUMPLFLN 40XW, Olympus). We took care to only record from GFP expressing cells. Electrophysiological signals were amplified and low pass-filtered at 10 kHz using a MultiClamp 700B amplifier (Molecular Devices). These amplifier settings are not optimal––the frequency of the low-pass filter should have been set to be less than half that of the sampling rate––and thus frequencies above 5 kHz are aliased in our recordings. However, a power spectral density analysis of these recordings, and additional control recordings, revealed no significant power in the electrical signals from PFNa cells above 1-2 kHz, minimizing any aliasing-related concerns. We streamed all voltage and current signals continuously at 10kHz with a Digidata 1440A input-output board (Molecular Devices). All data presented are *Vm* measurements in current clamp mode. We injected a small amount of current (-1 to -3 pA) during *Vm* recordings, to partially address the depolarizing effect of the seal conductance on small cells, with high input resistance. Out of 41 PFNa cell recordings, across all genotypes, in which we were able to present the entirety of the airflow-direction protocol, we excluded one cell from analysis because the fly did not walk enough to allow us to estimate a heading tuning curve. We excluded two additional cells because their baseline membrane potentials were above our threshold *Vm* for analysis, which was –40 mV before liquid-liquid junction potential correction, or, equivalently, –53 mV after correction. *Vm* measurements reported in the paper are corrected for a –13 mV liquid-liquid junction potential. Details on the passive membrane properties for each recording are listed in **Table S2**.

#### Closed-loop visual environment

We presented visual stimuli on a panoramic LED array^81^ spanning 270° in azimuth and 81° in height, with ∼1.875° pixel resolution^24^. The arena consisted of blue LEDs (BM-10B88MD, Betlux Electronics), covered by 7 sheets of blue gels (Tokyo Blue, Rosco) to reduce detection of light from the display by the microscope’s photomultiplier tubes. In patch-clamp electrophysiology experiments, we used the same LED display with 6 sheets of blue gels, 3 sheets of Clear-Shield film (Less EMF A1210-24) (that were electrically grounded) and one layer of stainless steel wire cloth (McMaster-Carr 85385T89) to minimize optical reflections between opposite sides of the display. In all experiments, we typically presented on the LED arena a 6-pixel (11°) wide bright blue vertical bar against a dark background. The bar rotated in closed loop with the fly’s yaw turns on an air-supported ball, thus simulating the movements of a distant landmark that can be used for orienting^24^. The rotation of the air-supported ball along the pitch, yaw and roll axes was detected via image analysis, using a custom-modified version of the open-source software FicTrac 2.0^82^. We modified the FicTrac 2.0 Python code to implement it within the Robot Operating System (ROS) platform, running it at 50 Hz during electrophysiology experiments and 80 Hz during imaging experiments. We illuminated the ball using 850 nm LEDs (OSRAM Platinum DRAGON, SFH 4235) guided by optical fibers (02-535, Edmund Optics). We imaged the ball using a Chameleon3 camera (CM3-U3-13Y3M, Teledyne FLIR) with an InfiniStix lens (144100, Infinity Photo-Optical). The lens was equipped with an 850/50 bandpass filter (84-778, Edmund Optics).

#### Design and construction of the airflow delivery device

We designed an airflow delivery device based on a previously published apparatus^59^. The previous apparatus employed a single airflow outlet pointing at the fly, which could rotate to any arbitrary angle around the yaw axis. We modified this device to ensure that nothing visual changed in the field of view of the fly when the airflow direction needed to be altered. Our goal was to make it possible to present separable visual and airflow stimuli to the fly; with the original device, whenever one would change the airflow direction, the rotating nozzle necessarily created a concomitant moving visual stimulus. To solve these potential stimuli confounds, the new apparatus in this study separated the rotating outlet from the airflow delivery part, which took the form of a static disc surrounding the fly. The rotating outlet in the new device is a circular assembly consisting of a stepper motor driving a rotating nozzle. The nozzle delivers air, whose flow rate is regulated by a mass flow controller, to a subset of 36 circularly arrayed airflow channels. Depending on the stepper motor’s position, different airflow channels are engaged. Each of these 36 airflow channels was connected to a matching airflow channel in the airflow delivery disc that surrounded the fly via flexible plastic tubing, this is the part of the device featured in the figure schematics throughout the paper. The static disc consisted of 36 tubular airflow channels, spaced evenly at 10° increments, with each channel pointing at the fly from a different angle around the yaw axis. The connecting plastic tubes attached to the outer rim of the disc were gathered using soft Velcro tape and funneled away from the fly’s field of view. In summary, a motor-controlled rotating outlet induced airflow into a distributor manifold connected to the tubes, which passed through specific airflow channels in the disc and ultimately hit the fly’s body from a specific direction around the yaw axis.

The airflow disc surrounding the fly was sufficiently thin in the vertical dimension that the fly could still see the vertical blue bar on the LED display. Thus, we could change the airflow direction and visual experience of the fly completely independently. The ∼1 m length of the plastic tubes does not significantly delay an airflow pulse from hitting the fly because air pressure changes can be considered as being transmitted instantaneously from one end of the tube to the other, for our purposes. Specifically, assuming a meter-long traveling distance, a airflow pulse from the rotating outlet would take ∼2.9 ms to travel across the tube, governed by the speed of sound.

We should note that, in its current incarnation, this device has two main disadvantages. First, it is not well suited for rapidly switching between different odorants, because unlike with air pressure changes, odor molecules need to travel down the entire length of each plastic tube before reaching the fly; this process would take much longer than a few milliseconds. Second, the airflow direction is discretized by the presence of the 36 static tubes, introducing variation in airflow speed depending on the phase of the rotating outlet relative to the holes in the first static ring. Specifically, we found that when we made the nozzle opening exactly the diameter of one airflow channel, the air speed arriving at the fly varied up to ∼50% depending on whether the nozzle was perfectly aligned with the channel or centered on the plastic midline between two channels, for example (data not shown). To combat this effect, we made the nozzle opening span two airflow channels, i.e., we made it 20° wide, which kept the proportion of the nozzle engaged with open air (within a channel) versus plastic more consistent independent of the nozzle’s position around the ring. This design feature minimized pressure changes that were contingent on the exact angle of airflow being delivered to the fly, while still maintaining a reasonable directional accuracy for the air stream.

##### 3D printing, part processing, and apparatus assembly

All custom parts were fabricated via 3D printing using VisiJet M3 Crystal material on a Projet MJP 3600 series 3D printer (3D Systems) at 30 µm resolution. We cleaned the printed parts using a multi-step procedure. First, the parts were incubated in an oven set to 65° C to melt and remove most of the wax support material. Afterward, the parts were sonicated in a mineral oil bath for at least one hour to dislodge the finer wax coating still attached to the plastic. The mineral oil residue was then removed by passing a jet of compressed air through all of the holes in the part. Afterwards, the parts were washed using water and dish soap (Dawn), rinsed with distilled water, and dried with compressed air. Each of the holes in the static airflow channels were manually tapped and fitted with a 1/16’’ brass hose barb (Clippard 12841) and the barbs on both parts (the airflow disc and the discretizing manifold on top of the rotating nozzle) were connected using soft plastic tubing (Tygon E-3603 via McMaster Carr, 5155T12). The rotating outlet component of the device was attached to an aluminum post and mounted on the same air table as the microscope. The airflow disc was mounted to the device that held the air-supported ball, which helped to ensure that the disc’s center was precisely located at the position in which the fly stood. Additionally, the airflow disc featured notches that nested the fly plates and ensured correct alignment of the fly’s body relative to the airflow disc during each experiment.

##### Airflow calibration

We calibrated the device by measuring the air speed at the center of the airflow disc with a hot wire anemometer (Climomaster Anemometer 6501-CE, Kanomax) fitted with an omnidirectional spherical probe (6543-2G, Kanomax). Using custom components, specially printed for alignment purposes, we positioned the spherical probe in the precise position that a fly would occupy during an experiment. We then manually rotated the nozzle so as to align it with the frontmost (0°) airflow channel. At this zero position, the 20° nozzle thus fully spanned the central airflow channel as well as ∼5° each of the channels on either side of the frontmost channel. The airflow angles used in this paper were all multiples of 10° from this zero position, meaning that for all airflow stimuli we expect there to have been the same phase relationship between the nozzle opening and the airflow channels downstream of the nozzle. This fact makes it more likely that the air speed that the fly experienced was consistent across different delivery angles (because we noted changes in air speed that could occurred as a function of the phase relationship of the nozzle and the downstream, discrete, set of tubes). We experimentally verified that the air speed was consistent at the position of the fly, with the exact air puff angles used in the paper, using the anemometer. We also validated the directionality of the air puffs delivered to the fly by visualizing CO_2_ gas (dry ice) with a laser sheet. We performed the dry ice visualization in a separate but nominally identical assembly.

#### Simultaneous control of the airflow and bar stimuli

We used analog voltages––generated by the microcontroller that controlled the visual pattern on the LED display^81^––to control the angular position of the airflow stimulus as well as the mass flow controller that turned the airflow on and off. To control the angle of the airflow stimulus, the analog voltage controlled the position of a stepper motor that drove the rotating air outlet to new set points via a custom-made analog-to-digital controller box. To control the air speed, a voltage signal controlled the opening and closing of 2-SLPM mass flow controller (Alicat Scientific). The approach of having the LED-arena’s microcontroller also drive the airflow stimuli made onset and offset latencies of the air puffs more reliable and also aided temporal alignment between the visual and air-puff stimuli.

#### Experimental protocol for presenting open-loop airflow and closed-loop visual stimuli

To solidify the visually mapping between the position of the blue bar on the LED display and the EPG bump position in the brain, all electrophysiology experiments and most imaging experiments began with a period of 5-10 minutes of the fly interacting with the bright blue bar in closed loop. This procedure was particularly important for electrophysiological data collection because we could not directly estimate the EPG phase in these experiments, and thus any changes in the mapping between the EPG phase and the visual cue would have manifested as multi-peaked heading tuning curves from single neurons. We could be less strict on this matter in two-photon imaging experiments because we were directly monitoring the EPG neuron population and could always estimate its phase in the brain, independent of the bar’s position on the LED arena. Following the acclimation period for establishing a robust EPG visual mapping, we began to present air puffs in open loop. We presented air puffs from twelve angles around the fly, in 30° increments, to fully and equally cover 360° of azimuthal space. The 0° direction, where air arrived from directly in front of the fly, was always included, which anchored the eleven other angles presented. Each open-loop air pulse lasted for 4 s with a 5 s inter-pulse interval. The air-puff angles were pseudorandomized in each block of twelve trials. We collected 3-5 blocks of trials per fly during imaging experiments, and 10 blocks during electrophysiology experiments. We used an airflow speed of 20 cm/s for all puffs. In this study, 0° represents the direction directly in front of the fly, and 180° represents the direction directly behind the fly. Negative angles represent air arriving to the fly’s egocentric left side, and positive angles other than 0° (front) and 180° (back) are to the egocentric right of the fly.

#### Synchronized acquisition of behavior and physiology data

We recorded all experiment-related signals in the form of voltages using a Digidata 1440A I/O board and Axoscope software (Molecular Devices). The air-supported ball’s yaw, pitch, and roll angles, the control signals to and from the airflow stepper motor, the Alicat flow meter output, the angular position of the blue bar the LED display, the internal triggers of the FicTrac ball-tracking camera, and custom signals to identify trials epochs, were recorded at 10 kHz in all experiments. Signals specific to either calcium imaging or electrophysiology, such as the membrane potential, injected current, or triggers for two-photon imaging frame acquisition, were recorded in addition when applicable. Behavioral, electrophysiological, and calcium imaging signals updated at different rates and we used either the FicTrac camera’s triggers, or the two-photon imaging frame triggers, for careful temporal alignments to behavioral and brain-imaging signals, as required.

#### Calcium imaging data analysis

##### Data processing

All calcium imaging analysis was performed using custom scripts written in Python 3.7 and the NumPy, SciPy, pandas, matplotlib, and seaborn libraries.^83–89^ We registered the raw time series of fluorescence images using either the rigid motion correction algorithm in the CaImAn software suite^90^ or a custom algorithm^24^ where shifts were applied to each frame in *x* and *y* to match a time-averaged frame for each *z*-plane. Both registration approaches produced qualitatively similar results. When available, we used the tdTomato or jRGECO1a image for registration, otherwise, we used the jGCaMP7f signal directly. After registration, we drew regions of interest (ROIs) manually, in each z-plane separately, using custom software^80^ written in Python 3.7.^83^ We drew ROIs on fluorescence images averaged across an entire recording session, and our assignment of glomerular boundaries was aided by simultaneously viewing autocorrelation images using data from the entire session. When analyzing data in the protocerebral bridge, individual glomerulus identities were assigned by adhering to the following established anatomical principles^63^: (1) the entire protocerebral bridge should be composed of 18 distinct glomeruli, (2) PFN neurons do not innervate the pair of glomeruli closest to the midline, and (3) EPG neurons do not innervate the pair of glomeruli furthest from the midline. Out of a total of 12 collected bridge co-imaging datasets, we did not analyze 4 recordings in which more than one glomerulus was not visible, or whose identity could not be clearly established. When analyzing the ellipsoid body, we divided the structure into 16 evenly spaced radial sectors, whose boundaries and center were defined across every plane^11,24^. Following previous conventions^11,24^, and to establish a consistent anatomical correspondence with the bridge and fan-shaped body, wedge 1 and wedge 16 of the ellipsoid body were defined, respectively, as the wedges immediately to the left and immediately to the right of the vertical bisector line at the bottom of the torus, when viewed from the posterior side of the head. When analyzing the fan-shaped body, we defined columns as follows. We first drew an outline of the entire structure. Then, the left and right edges of the structure were marked with angled lines. The vertex point at which the extension of these two angled lines met defined the entire angular extent of the fan-shaped body, which we then subdivided into 16 equally-spaced columns^11^. Column 1 of the fan-shaped body, which corresponds functionally to wedge 1 of the ellipsoid body, was defined as the leftmost fan-shaped body column when viewing the structure from the posterior side of the head. Noduli outlines were manually drawn and assigned as either left or right, using the same posterior view.

##### Fluorescence signal normalization

For each imaging volume and structure being imaged, we averaged together the signal from ROIs across z-planes, if they belonged to the same glomerulus, wedge, column or nodulus side, thus generating a final, unidimensional array of fluorescence intensity values for each structure at each time point. Each of these unidimensional arrays, corresponding to individual sector ROIs, were concatenated into a matrix where each data column corresponded to one sector and each row corresponded to a time point. Thus, the matrices of fluorescence values were composed of 18 data columns for the 18 glomeruli in the protocerebral bridge, 16 data columns for the 16 wedges of the ellipsoid body, 16 data columns for the 16 anatomical columns of the fan-shaped body, and 2 data columns for the 2 sides of the noduli. We report all imaging data from the ellipsoid body and fan-shaped body as ΔF/F, calculated as (F – F_min_) / (F_min_). In this equation, F corresponds to the raw fluorescence intensity measured in a given sector ROI (i.e. in each wedge or column) and F_min_ corresponds to the 5^th^ percentile of F values observed in that same sector ROI over the whole recording. When analyzing calcium imaging data from the protocerebral bridge, we normalized each glomerulus ROI to its own maximum and minimum F values because some glomeruli were much brighter or dimmer than others likely due to the expression levels of the fluorophore. Thus, the values of ΔF/F_max_-F_0_ that we report here for the protocerebral bridge correspond to (F – F_min_) / (F_max_ – F_min_), where F corresponds to the raw fluorescence signal in sector ROI, and F_min_ and F_max_ represent the 5^th^ and 95^th^ percentile of the F values from that same sector ROI over the entire recording. When analyzing the data from the noduli, we performed a z-score normalization over the raw fluorescence values, calculated separately for each of the left and right nodulus sector ROIs.

##### Aligning fluorescence readings with behavioral measurements

We matched behavioral (80 Hz) and imaging (4-8 Hz) measurements in the following manner. We determined the epochs corresponding to entire volumetric scan cycles using the voltage triggers output by the two-photon microscope, and we averaged the fluorescence values corresponding to each ROI across all z-slices containing them in the cycle. We averaged behavioral-related measurements, such as the visual stimulus’ position on the screen, over the time corresponding to an entire volumetric scan cycle as well. The behavioral signals are thus averaged over epochs of ∼125-250 ms (corresponding to the imaging period given our scanning rates of 4-8 Hz).

##### Phase extraction

Columnar neurons in the central complex often express spatially localized calcium signals that are referred to as bumps of activity. Because space around the fly, 0°-360°, physically maps to positions in central complex structures, the position of the calcium bumps within a structure has an angular interpretation and thus is often referred to as the bump’s phase. We extracted the phase of calcium bumps in the ellipsoid body and fan-shaped body by taking a population vector average (PVA)^15,24,26^. We computed the PVA by taking the circular mean of the calcium signals across all ROIs associated with fixed angular positions, corresponding to each wedge in the ellipsoid body or glomerulus in the protocerebral bridge. The “PVA strength” measure corresponds to the length of the resultant vector generated when taking the circular mean. We defined the sectors of the ellipsoid body and fan-shaped body to have their numerical values (1 to 16) match the known anatomical and functional correspondence across these two structures. Thus, when phase of bumps in these two structures matched, this meant that the angular signal carried by the two neuronal populations were functionally aligned in angular space, and phase differences across the ellipsoid body and fan-shaped body could be calculated through a simple angular subtraction as described in the *Data processing* subsection above. When imaging EPG neurons in the protocerebral bridge, the phase of the EPG heading estimate was extracted as previously described^24^. In brief, we took a Fourier transform of the sector ROI signal over the 16 glomeruli. The phase of the EPG bump in a given time point was then defined as the phase of the Fourier component with a wavelength of 8 glomeruli. This approach assumes that an invariable 8-glomerulus spacing exists between the peaks of the two EPG bumps in the bridge, and that, together, these two bumps encode a single shared heading angle estimate, which has proven to be a robust assumption.

##### Phase nulling

To focus on the relationship between the EPG and PFNa phases, we computationally shifted the EPG bumps in each imaging frame to the same angular position in the bridge and rotated the PFNa bumps by this same EPG-determined angle. This *phase-nulling* analysis^11,24^ helps to visualize the alignment between the EPG and PFNa bumps by removing the complexity of the bumps moving around the bridge. We phase-aligned EPG and PFN bumps in the protocerebral bridge using an algorithm described previously^24^. In brief, the sector ROI arrays from the protocerebral bridge for EPG and PFNa cells were first, separately, interpolated to 1/10th of a glomerulus resolution using a cubic spline. We then circularly rotated the data array at every time point so that the EPG phase was at the same position along the x axis. This EPG-determined rotation was then applied to the PFNa interpolated vector signal, allowing us to visualize the phase of the PFNa bumps in reference to that in the EPG system. It is only by employing this phase nulling analysis that we analyzed the phase of the PFNa bumps presented in this study. We found that we could not easily define the phase of the PFNa bumps otherwise, because they were not stably periodic nor always visible in calcium experiments.

##### Estimation of the phase-aligned and phase-inverted response amplitudes

We applied a constrained non-linear optimization function (fmincon in MATLAB) to the phase-nulled calcium imaging data of the PFNa neurons to determine the amplitudes of the phase-aligned and phase-inverted responses to airflow. To do this, we found the best sinusoidal fit for the phase-nulled responses (phase of the phase-nulled response equal to 0) and phase inverted response (phase of the phase-nulled response equal to 180), separately, across all airflow angles. The best-fit amplitudes are reported in Figures 2E and 2F.

#### Electrophysiology data analysis

All electrophysiology analysis was done using custom scripts written in Python 3.8.^83^ We processed current-clamp recordings to extract each cell’s membrane potential (*Vm*), sodium-spike rate and 2-6 Hz power (due to calcium spikes).

##### Sodium spike detection and processing

To detect sodium spikes, we bandpass-filtered the raw 10 kHz *Vm* traces in the range of 300-1000 Hz using an 8-pole Bessel filter. Sodium spikes appeared as large deflections in this high-passed filtered signal. Second, we isolated discrete moments in the bandpass filtered trace in which a positive-sloped crossing of a high threshold occurred. The threshold values ranged from 0.075 to 0.28 mV; these values were selected depending on the signal quality of each recording. Threshold crossings were candidate moments in which a sodium spike was fired. To finalize a spike-event call, we used one of two different approaches. In one approach, we required that the mean *Vm* of the cell at the moment of threshold crossing exceed a *Vm* threshold, to exclude potential artifactual events that occurred much below a *Vm* were a sodium spike is expected. We rejected < 1% (∼0.9%) of threshold crossing events (average across 20 cells) based on this final *Vm* threshold criterion. In the second approach, we called all threshold crossing events spikes. This second approach seemed sensible because spikes could occur at the top of calcium spikes, where the somatic *Vm* is strongly influenced by T-type currents, and sometimes our own current injection, which means that the somatic *Vm* is likely to be quite different from the *Vm* at the spike initiation zone. In other words, we should be open to spikes occurring at *Vm* levels that seem more hyperpolarized than usual. We used this second method––of not rejecting spikes below a *Vm* threshold––when analyzing sodium spikes occurring at the peak of calcium spikes and during our analyses of spike rate during current injection. Regardless of the approach used, the difference between these two algorithms for calling spikes should impact only <1% of spike assignments. For each recording and cell, we visually inspected a subset of the events to verify that appropriate events were being called spikes.

We converted sodium-spike times to spike rate by convolving spike times with a sliding 1-second-wide Hann window. Sodium spike amplitudes were often small––only a few millivolts or less––in these recordings, which is typical of recordings from somata of neurons in *Drosophila*. As a result, any spike-detection algorithm used is likely to have resulted in calling some non-spike events as spikes and to not have detected some number of genuine sodium spikes emitted by the cells. However, our results depend on an overall estimate of spike rate and not precise spike times; some low level of spike misdetections might raise or lower the estimate rate over time, but this should not drastically impact the shape of the relevant tuning curves for our model.

##### Calcium spike quantification

We quantified calcium spikes in two ways. The main quantification employed in this study was the membrane potential oscillation strength in the 2-6 Hz range. To extract oscillation strength, we downsampled raw *Vm* traces to 100 Hz and performed a fast Fourier transform on the traces (specgram function in Matplotlib; window size = 4 s; window step size = 20 ms). We averaged the calculated power of *Vm* frequencies in the range of 2-6 Hz and used this value as our metric for estimating the strength of T-type oscillations in PFNa neurons. To quantify the oscillation strength during 2 s periods of current injection, we first conditioned the current-injection *Vm* traces as follows. We started by defining two 600-ms time segments. One segment started 550 ms before the start of the current injection pulse and the second segment started 50 ms before the end of the current injection pulse. We then replaced the actual *Vm* sample points in both segments with a linear interpolation between the first and last sample point within the time segment. We then took the fast Fourier transform of the *Vm* traces and quantified the power in the 2-6 Hz band, using a window size of 2 s and window step size of 0.01 s. The linear interpolation of the *Vm* trace that we performed at the onset and offset of the current pulse prevented artifactual frequencies from appearing to have high power in the Fourier transform analysis simply due to the large *Vm* step-like changes that necessarily occur at the start and end of the current injection pulse. This quantification allowed us to detect oscillatory activity at the more depolarized *Vm* ranges, where it was difficult to distinguish individual calcium spikes due to their low amplitude. In the second quantification, we detected calcium spikes directly. To do so, we low-pass filtered the *Vm* signal with a cutoff of 100 Hz, and detected peaks of determined prominences (typically 3-5 mV) and widths (typically 50-150 ms) in the filtered *Vm*. This detection algorithm performed best at the more hyperpolarized baseline *Vm* values for two reasons. First, the calcium spikes had a larger amplitude when the membrane was substantially hyperpolarized, and thus the spikes were more prominent. Second, the background subthreshold activity of the cell was lower when the cell was more hyperpolarized, and thus the algorithm suffered from fewer instances of mis-classification of a subthreshold event (such as an excitatory post-synaptic potential) as a calcium spike. We classified calcium spikes as containing a sodium spike at the peak if a sodium spike was detected within ±50 ms of the peak of the calcium spike.

##### Processing of the *Vm* signal, resampling, and data averaging

Raw *Vm* signals were always corrected for a 13 mV liquid-liquid junction potential. For summary analyses where we averaged waveforms across multiple cells, we downsampled the 10 kHz *Vm* traces to 1 kHz. To compute average *Vm* values during airflow, we removed spikes from the trace by applying a median filter using a kernel size of 40 ms. To align behavioral readings with electrophysiological measurements, the sample points from the FicTrac camera –running at 50 Hz in patch-clamp experiments– were upsampled to 1 kHz using linear interpolation. This upsampling of the behavior-related signals could then be directly aligned to the analyzed *Vm* signal, which was downsampled to 1 kHz as described above.

##### Estimation of baseline *Vm*

PFNa neurons express large calcium spikes, alongside large synaptic potentials, even when very hyperpolarized, which makes defining a resting *Vm* difficult for these cells. Thus, instead of estimating a resting *Vm* for PFNa cells, we instead defined the baseline *Vm* as the minimum value in the heading tuning curve calculated for each cell (mean: -64 mV, standard deviation: 4.8 mV, range: 19.9 mV). This baseline *Vm* value was typically stable during our recordings, which usually lasted 45 to 60 minutes. Whereas the baseline membrane potential varied substantively across cells (**Figures S3A and S3B**), we found that the peak-to-minimum membrane potential amplitude of the heading tuning curve (mean: 13.7 mV, standard deviation: 2.9 mV), and the sodium spike rate at the minimum value of the heading tuning curve (mean: 0 spikes/s, standard deviation: 0.1 spikes/s) was more consistent across cells. Given this observation, membrane potential data was combined across cells by subtracting the baseline membrane potential from each cell. We compare tuning curves with and without normalization in **Figure S3**. Spike rate data were combined across cells without any such normalization.

##### Estimation of heading, airflow direction and conjunctive tuning curves

For each PFNa cell, we generated *Vm*, spike rate, and oscillation strength tuning curves as a function of heading and airflow angle. We generated single-variable tuning curves against these two variables, and we also generated conjunctive, two-variable (heat map) tuning curves against the two variables. For heading tuning curves, we binned the time series of bar positions on the LED display into 10° bins, and we averaged the *Vm*, spike rate or oscillation strength in each bin. We required that a bin included at least 2 s worth of sample points for its average to be calculated. We also required that the fly not be standing (forward speed > 0.5 mm/s) for sample points to be included in the tuning curve. We calculated the preferred heading direction of a given cell’s tuning curve by finding the angular shift of the curve that produced the highest correlation between the actual tuning curve and a normalized cosine function. We calculated tuning curves to air puffs by averaging the *Vm*, spike rate, or oscillation strength in the 10 air-puff trials for each of 12 air puff directions presented in a 2 s window, starting 0.5 s after the onset of the air puff. To estimate conjunctive tuning to heading and airflow, we split the data into 18 (20° heading) x 12 (45° airflow direction) bins. We required that at least 0.5 s of data populate each bin in order to calculate a mean value. After calculating the conjunctive tuning heat-map for each cell separately, we generated combined heat maps across cells after centering each single-cell heat map on the preferred heading direction of a given cell.

#### Model construction

Membrane potential data over 12 egocentric airflow directions, *W*, and 18 relative heading directions, *H* (the heading direction relative to the preferred heading direction for a particular neuron), averaged over individual recorded PFNa left and right neurons, were fit using the form *V_m_* = *a_o_* + *a_H_*cos(*H* ) + *a_W_* cos(*W* ± *n* /4), with free parameters *a_o_*, *a_H_* and *a_W_*, and the ± applied to left or right neurons, respectively. This 3-parameter fit of 216 data points for each case (left/right) explained 92% (left) and 94% (right) of the variance of the data.

Firing rates for sodium spikes were fit to the equation *r* = ([*b_H_*cos(*H* ) + *b_W_* cos(*W* ± *n* /4)] )^2^, with *b_H_* and *b_W_* free parameters and [*x*]_+_ = *x* for *x* > 0 and 0 otherwise. This two-parameter fit of 216 data points (as above) explained 91.1% and 87.2% of the variance for the left/right PFNa neurons, respectively. We fit the oscillation power for these data to *p* = *c*_0_ + ([*c_H_*cos(*H* ) + *c_W_* cos(*W* ± *n* /4)]_)^2^, with *c*_0_, *c_W_*, and *c_H_* free parameters and [*x*]*_−_* = *x* for *x* < 0 and 0 otherwise. This three-parameter fit of 216 data points explained 84.4% and 70.1% of the variance for the left/right PFNa neurons.

We considered various sums of the plus and minus rectification terms discuss above, parameterized by the relative strength of the negative rectification term. We also considered the case when the plus and minus rectifications have equal strength such that the total output can be expressed as (*b_H_*cos(*H* ) + *b_W_* cos(*W* ± *n* /4))^2^, with no rectification as a consequence of summing sodium and calcium spike contribution. In this case, summing the left and right contributions and accounting for the ±45° anatomical shifts of the PFNa projections from the protocerebral bridge to the fan-shaped body, as well as the fact that airflow input to the noduli comes from the opposite side of the midline from the heading input, gives the total signal from this similarly tuned pair as

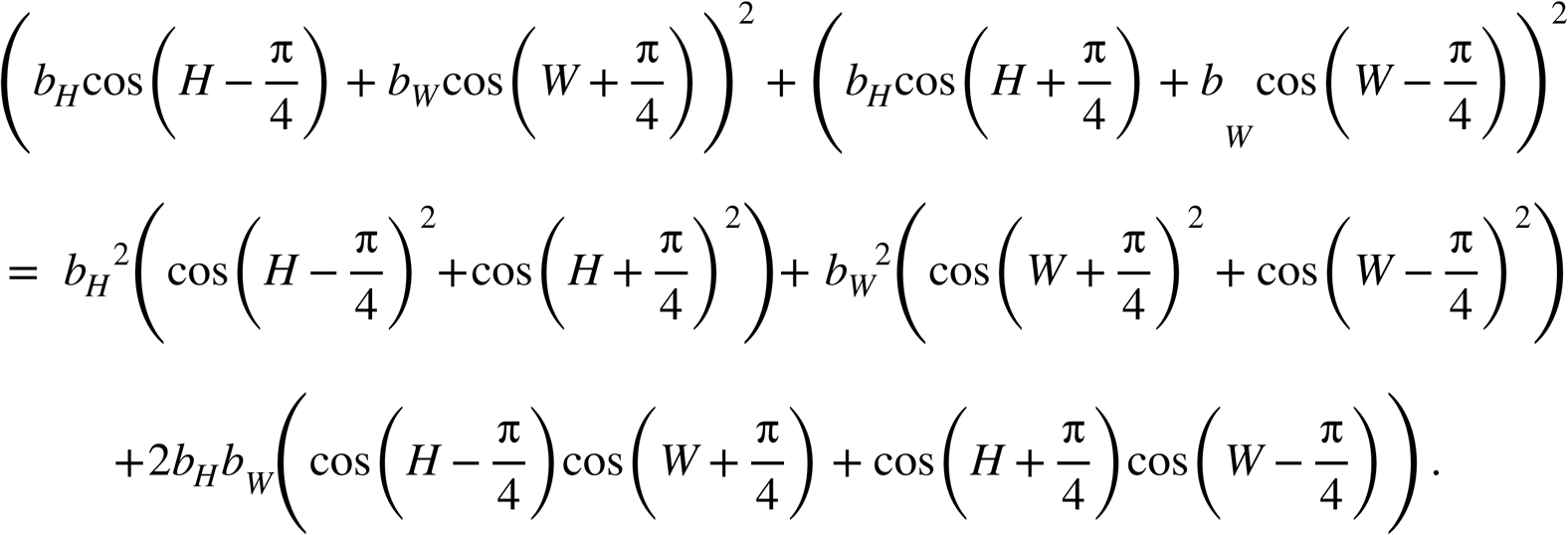

Because cos(*x − 1r* /4) = sin(*x* + *1r* /4), and the sum of squared sines and cosines is 1, the first line on the right of the equal sign above equals *b_H_*^2^ + *b_W_* ^2^. Using the sum-of-angles identity, cos(*H + n* /4)cos(*W* ± *n* /4) = _(_cos(*W* + *H*_)_ + cos(*W − H* ± *1r* /2))/2, and noting that cos(*x* + *1r* /2) + cos(*x − 1r* /2) = 0, the last expression above becomes 2*b_H_b_W_* cos(*W* + *H* ). Putting this all together, we find that the total output of this pair of PFNa neurons is *b_H_*^2^ + *b_W_* ^2^ + 2*b_H_b_W_* cos(*W* + *H* ). The output across the full population of PFNa neurons, parameterized by their preferred heading angle is then a phasor representing a vector of unit length pointing in the allocentric direction of the airflow. Note that this result requires that we sum the contributions of left and right PFNa cells, represented by the two terms in the first line of the above equation. If we consider the left and right PFNa signals separately, we obtain

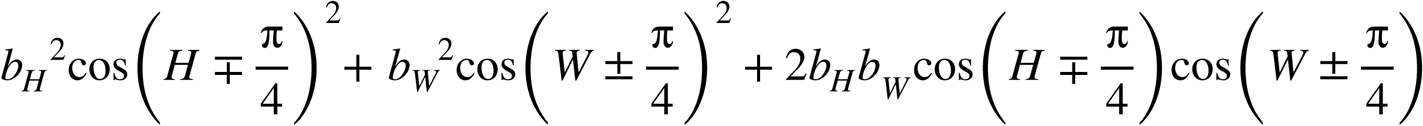

for the two sides. When extended to the full left or right PFNa population, the last term represents the inverting vectors discussed through this paper.

At some airflow angles, all of the model PFNs neurons on a given side (left or right) are in the sodium spiking mode, and for other airflow angles, they all exhibit calcium spikes. However, at intermediate angles, some of these neurons can fire sodium spikes and others can fire calcium spikes. In this case, the model predicts that two bumps, offset by 180° should be visible in the protocerebral bridge. Indeed, such an effect can be seen in the data (**Figure S2**, airflow angles ±120° and ±150°). Mathematically, this is due to extra terms in the PFNa responses other than those that encode the left and right component vectors. Because these additional terms cancel when the left and right PFNa response are summed (and thus they have no effect on the airflow vector encoding), and because their impact is only seen in the calcium data when the PFNa signal is weak (**Figure S2**), we have not emphasized them in our discussion.

##### Estimation of the encoded angle in the PFNa neurons’ calcium signals

We used the phase-nulled responses of the PFNa neurons (data in **Figure S2A**) to estimate the encoded output angle across the population. Because the GCaMP fluorescence intensity of the phase-aligned and the phase-inverted responses is different –likely due to the distinct biophysical origin of each– we introduced a correction factor to normalize both types of responses to a common range. Specifically, we applied a gain factor of 0.2 to the bridge glomeruli that display phase-inverted responses. The gain factor was chosen by optimizing the match between the decoded airflow direction and the actual airflow direction. The glomeruli at which to apply the gain factor were determined by inspecting the bridge responses at the preferred airflow directions for eliciting calcium spikes, ±120° in our experiments; the assumption is that the phase-inverted responses correspond to glomeruli whose constituent neurons express calcium spikes. There are some cases where assigning whether a particular PB glomerulus as sodium- or calcium-spiking is ambiguous, i.e. the glomeruli that lie in-between the phase-aligned and the phase-inverted bumps. However, these glomeruli contribute little to the decoding because their responses are nearly flat. As a result, if we vary the gain factor associated with the ambiguous glomeruli, it makes little difference in the encoded angle, suggesting that the exact assignment of the ambiguous glomeruli as phase-aligned or phase-inverted is not critical for this analysis.

##### Prediction of the FC3 phase using the electrophysiology data of PFNa neurons

We interpolated the PFNa spike-rate and oscillation power data in Figures 3H and 3I and shifted them along the heading direction axis to duplicate the shifts that occur in PFNa projections from the protocerebral bridge to the fan-shaped body. The angles used for these shifts were obtained by averaging the two forward (for spikes) and two backward (for oscillation power) angles in **Figure S7F**. We also normalized these data so that we could compare the strengths of the spiking and oscillation contributions. The normalization was performed by subtracting the minimum value of each dataset (spike-rate or oscillation power) and then dividing over the maximum value.

#### Analysis of the open-source hemibrain connectome dataset

All electron microscopy-based analysis in this study was performed using the hemibrain connectome^60^ dataset v1.2.1 and neuPrint-python^91^ v0.4.15. Skeleton renderings were performed using the NAVis^92^ python library v1.3.1.

#### Analysis of the vector projection axes

When we fit our model to *Vm*, spiking, and oscillation data, we used fixed angle offsets of ±45° for the airflow tuning curves. However, we also fit these data allowing these offset angles to be free parameters. When we did this for the *Vm* data, we obtained angles of -41.3° and 41.9°. Although these angles differ from the ±45° assumed in our model, the improvement in the fits obtained by introducing these extra free parameters was small, only 0.09% additional variance explained for the left-bridge and right-bridge PFNa neurons. Allowing the angles to be free parameters for fitting the spiking and oscillation data gave angles of -38.9°, 38.8°, -61.1°, and 51.6° (**Figure S7F**) for the left-bridge and right-bridge spiking data and left-bridge and right-bridge oscillation data, respectively. This allowed the fits to explain an additional 1.3%, 1.4%, 1% and 7.5% of the variance in the data. These angles are close to, but slightly offset from, the ±45° expected from a perfectly orthogonal system.

The deviations from orthogonality suggested by the above analysis make quantitative predictions about the nature of heading tuning in the PFNa neurons if a vector sum process aims to accurately signal the allocentric airflow direction. For example, if a PFNa cell were to express its maximal airflow response at +40°––i.e., 5°closer to the fly’s midline than the +45° orthogonal prediction––its response to heading in the fan-shaped body should be shifted by +50° (relative to its response in the protocerebral bridge), i.e., 5° further from the midline, for the vector sum process to be accurate. Put another way, if a PFNa-encoded vector expresses its maximal vector length when airflow arrives from +40°, then the axis that this vector points along should be +50° for the vector sum process to be accurate. With non-orthogonal axes for a pair of basis vectors, the angle of each axis and the airflow angle at which each basis vector has its maximal projection to that axis are no longer the same angle.

To determine the angular shift in the PFNa projection from the bridge to the fan-shaped body––which determines the axis along which each PFNa vector points––we used the hemibrain connectome^60^. In the bridge, we assigned an angle to each synapse that a PFNa cell receives from an EPG or Δ7 cell, based on standard assumptions of how heading is signaled in the EPG/Δ7 system^11,16^ (**Figure S7A**). We averaged all the synaptic angles onto a given PFNa cell to assign each PFNa cell an overall heading angle for which it codes. We then analyzed the pattern of PFNa synapses onto a specific recipient cell class in the fan-shaped body, FC3 neurons. We found that the mean angular deviation between the left- and right-bridge PFNa cells that co-innervate FC3 cells in a given fan-shaped body column is 90.5° if EPG cells were considered to be the only drivers of PFNa activity and 113° if Δ7 cells were considered to be the only drivers (**Figures S7A, S7B, S7C, and S7E**). Because EPG and Δ7 cells express 11% and 89% of the synapses onto PFNa cells, respectively, their circular weighted average predicts a deviation of 110.6° between the left and right PFNa vectors as they influence a given FC3 column (**Figures S7C and S7D**). This anatomical calculation thus argues that the projection axes for the two PFNa vectors are offset by 55.4° to the left and right of the midline (**Figure S7D**). As mentioned, the fact that this angle is greater than 45° is expected given that the peak vector lengths, assessed with physiology, were measured to be closer to the midline than 45°. Specifically, our physiological measurements of ±39° predicted projection axes of ±50-51°, which are in reasonable alignment with the 55° estimate extracted from the connectome. The fact that the FC3 phase rarely deviated by more than ±55° from the EPG phase in response to air puffs (**Figure S6E**) is consistent with the possibility of the anatomical projection axes of the two PFNa vectors being separated by ∼110°, as the connectome analysis suggests. This offset angle would limit the maximal deviation of any downstream bump that sums the two front vectors.

Together, our angular measurements across the anatomy and physiology of the PFNa neurons suggest that its vector system could function in a non-orthogonal manner (**Figure S7G**). However, given the rather modest improvements provided by the fitted angular variables mentioned in the first paragraph above, we chose to use the canonical angles of ±45° for our model.

#### Immunohistochemistry

We dissected adult fly brains and fixed them in 2% paraformaldehyde for 55 minutes in 24-well crystallization plates (Cryschem M Plate, Hampton Research). We washed the brains 3x for 20 minutes each in phosphate-buffered saline containing 0.5% Triton-X (PBST), then blocked them with 5% normal goat serum (NGS, sourced from Gibco). For immunostaining GFP, we used a primary antibody solution consisting of 1:1000 chicken anti-GFP (600-901-215, Rockland Immunochemicals),1:30 mouse anti-bruchpilot (nc82, Developmental Studies Hybridoma Bank), and 5% NGS diluted in PBST. For visualizing smFLAG-vGlut and mCherry expression, we used rabbit anti-dsRed (1:500, Takara Bio 632496), rat anti-FLAG (1:200, Novus Biologicals NBP1-06712), and mouse anti-bruchpilot (1:10, Developmental Studies Hybridoma Bank nc82). For visualizing 9xV5-vGAT and mCherry expression, we used rabbit anti-dsRed (1:500) and mouse anti-bruchpilot (1:10) in 5% NGS. We nutated the brain samples in primary antibody solution for 4 h at room temperature, followed by an overnight rocking incubation at 4°C. Afterwards, we washed the brains 3x for 20 minutes each in PBST, then incubated them in a secondary antibody solution. For secondary immunostaining of GFP, we used a solution composed of 1:800 goat anti-chicken Alexa Fluor 488 (A11039, ThermoFisher Scientific), 1:200 goat anti-mouse Alexa Fluor 594 (A11032, ThermoFisher Scientific), and 5% NGS in PBST. For 9xV5-vGAT and mCherry visualization, the secondary antibody solution was composed of of goat anti-rabbit Alexa Fluor 488 (1:400, Thermo Fisher A-11034) and goat anti-mouse Alexa Fluor 633 (1:400, Thermo Fisher A-21052). This experiment required an additional blocking step in 5% normal mouse serum (NMS) after washing the secondary antibodies, followed by and an incubation with mouse anti-V5 conjugated to DyLight-550 (Bio-Rad MCA1360D550GA) in 5% NMS. For smFLAG-vGlut and mCherry visualization, the secondary antibody solution was composed of goat anti-rabbit Alexa Fluor 488 (1:400), goat anti-rat Alexa Fluor 594 (1:400), and goat anti-mouse Alexa Fluor 633 (1:400). We nutated the brains in secondary antibody solution for 4 h at room temperature, then incubated them overnight at 4°C. Afterwards, we washed the brains 3x for 20 minutes each in PBST, performed a final wash in phosphate-buffered saline, and mounted the brains onto glass slides in 8 µl of FocusClear (CelExplorer) with the posterior side up (i.e. the posterior end of the brain faced the coverslip). We imaged the mounted brains using an upright Zeiss LSM 780 confocal microscope fitted with a 20x 0.8NA air objective (Plan-Apochromat 20x/0.8, Zeiss). Each stack of images consisted of 80-120 optical sections spaced ∼1 µm apart.

#### Analysis of open-source RNA sequencing dataset

The data in **Figure 4B** were obtained from an open-source RNA sequencing dataset by Davis et al, 2020^28^, available in the NIH’s Gene Expression Omnibus (GEO)^93,94^ under accession number GSE116969. The specific dataset shown here is GSE116969_dataTable4.genes_x_cells_TPM.coding_genes_QCpass, which reports the cell type-mean abundance of protein coding genes in quality control-passed samples. The sample labeled as “PFNa” in Figure 4B is PB_5, which corresponds to sequencing data from cells labeled by the split-Gal4 line SS02255. No additional processing was performed on the reported abundances.

### Quantification and statistical analysis

Details on statistical testing, as well as exact p-values, are presented in **Table S3**. Throughout the paper, we used a circular mean function implemented after Fisher and Lee, 1983^95^. We performed t-tests using the scipy.stats package^85^ and we performed Watson-Williams tests using the pycircstat package, a Python implementation of CircStat^96^, a toolbox for directional statistics in MATLAB.

### Additional resources

Code for data analysis and modeling, as well as experiment scripts and 3D models of the airflow device and associated custom control box, are available at https://github.com/MaimonLab/cx-vector-inversion.

---

**Figure S1.**
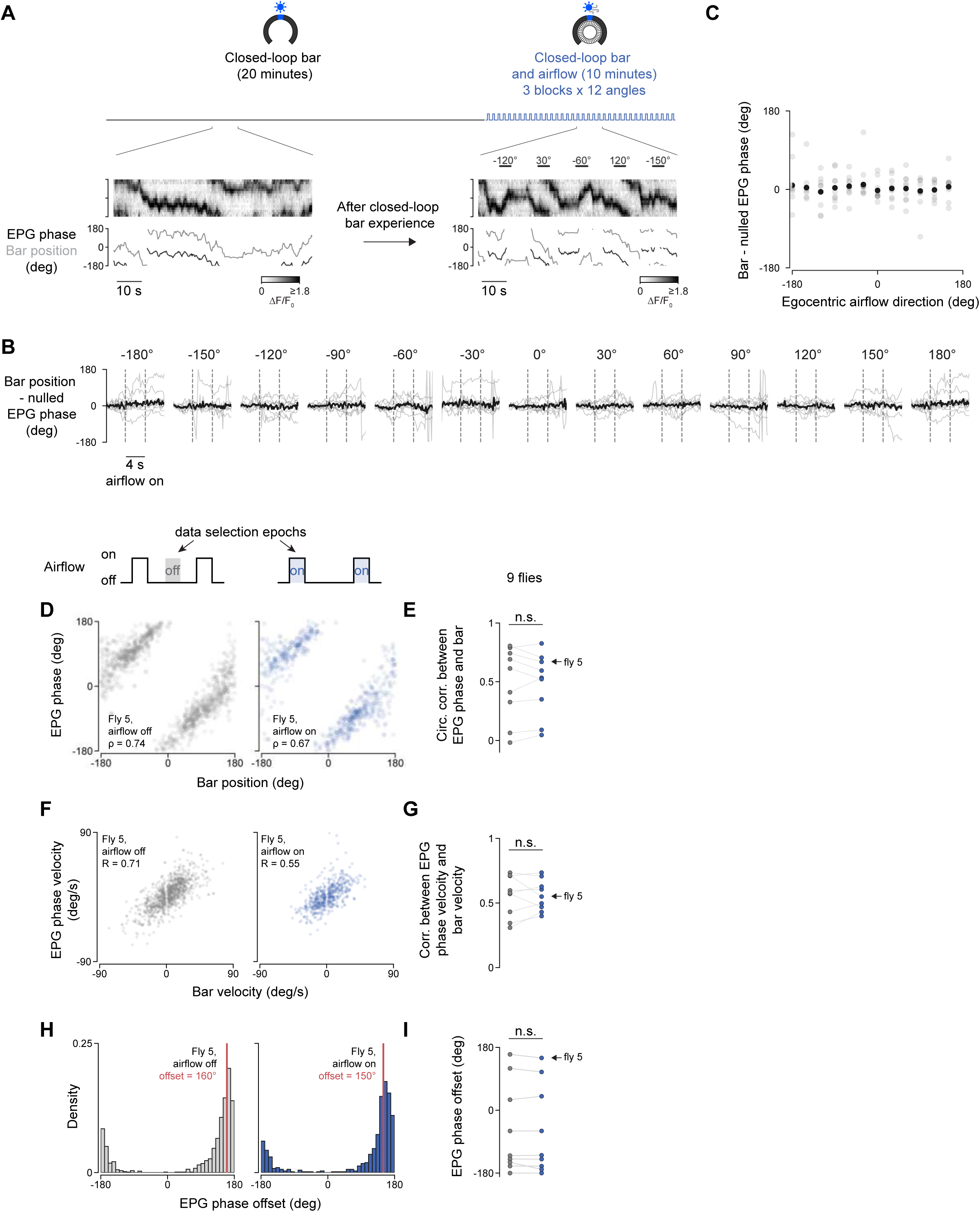
related to Figure 2. EPG neurons track the fly’s heading equally well in the presence and absence of open-loop airflow pulses. **(A)** _Example trace of EPG activity in the ellipsoid body in the closed-loop visual and open-loop airflow experiment. The ellipsoid body is split and unfolded as to display the donut as a single 16-wedge long array over time (gray color mesh), the color corresponds to the intensity of activity along each sector at every imaging frame. The position of the EPG activity peak is the population vector average estimate along the ellipsoid body, called the EPG phase (shown in black lines). The position of the visual stimulus on the LED screen is shown as a gray line. **(B)** Correspondence between the EPG phase and the bar position as a function of the 12 different air puff directions. The arbitrary offset between the EPG phase and bar position was zeroed at the beginning of each airflow trial, to highlight perturbations of the EPG phase-to-bar correspondence during the stimulus period. Average of 9 flies. **(C)** Average values of the last two seconds of airflow stimulation for the data shown in B. 9 flies. **(D)** Correspondence between the EPG phase position to the visual stimulus position on the screen in an example fly. The visual stimulus is never off: in the “airflow off” condition (left panel), only the closed-loop visual stimulus is present. In the “airflow on” condition (right panel), both the closed-loop visual stimulus and the open-loop airflow are present. The circular correlation coefficient p is noted. Data corresponding to moments where the fly stood still was excluded from this graph. **(E)** Circular correlation between the EPG phase and the bar position for 9 flies, with airflow on and airflow off. Gray dots show the “airflow off” condition, blue dots show the “airflow on” condition, and thin lines pair values for individual flies. The arrow highlights the data points corresponding to the example fly in D. **(F)** EPG phase velocity as a function of the visual stimulus velocity in an example fly. Pearson’s R is noted. Data corresponding to moments where the fly stood still was excluded from this graph. **(G)** Correlation between the EPG phase velocity and the visual stimulus velocity in 9 flies. Gray dots show the “airflow off” condition, blue dots show the “airflow on” condition, and thin lines pair values for individual flies. The arrow highlights the data points corresponding to the example fly in F. **(H)** Quantification of the arbitrary offset between the EPG phase and the visual stimulus for an example fly. The circular difference between the EPG phase and the bar position on the screen was divided into 36 bins of 10°, and the bin with the highest counts (labeled by the red line) was selected as the offset for the airflow on and airflow off conditions for each fly. Data corresponding to moments where the fly stood still was excluded from this graph. **(I)** Summary of the arbitrary offset between the EPG phase and the bar position on the screen for 9 flies. Each point in this plot represents the mean difference between the EPG phase and the bar position over the last 2 s of the airflow stimulus. Gray dots show the “airflow off” condition, blue dots show the “airflow on” condition, and thin lines pair values for individual flies. The arrow highlights the data points corresponding to the example fly in H._

**Figure S2.**
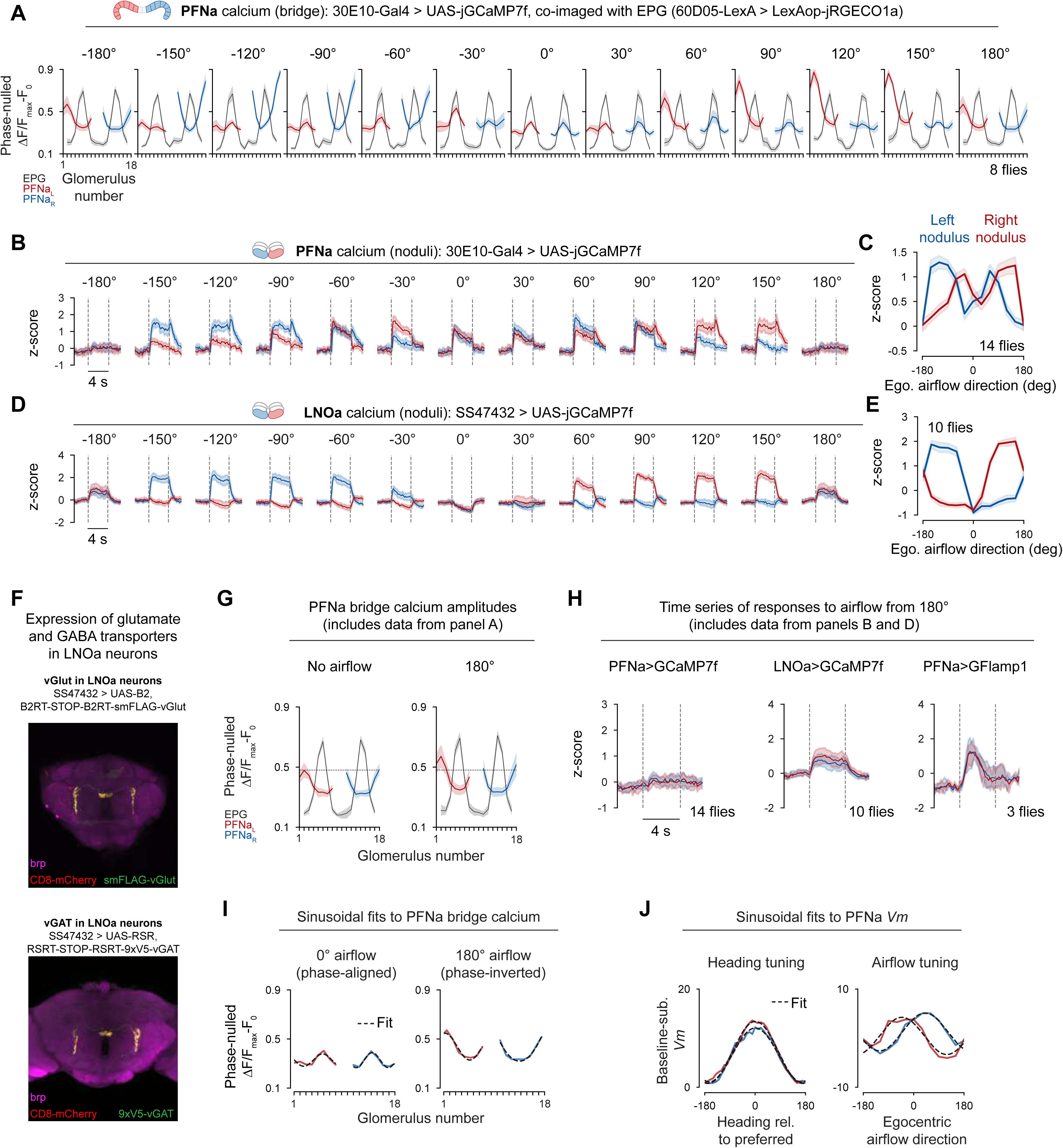
related to Figure 2. Tuning curves to airflow stimuli in the PFNa system. **(A)** Average data for the PFNa neurons’ phase alignment (red and blue traces) relative to the EPG heading neurons (black trace); we show the full series of airflow angles tested. We nulled the EPG phases on both sides of the bridge independently and at every imaging frame, corresponding to one time point. The EPG phase estimate at each time point was aligned to the middle glomeruli in a virtual protocerebral bridge. We then shifted the PFNa phase using the EPG phase estimate to determine its peak relative to the EPG neurons’ phase. **(B)** Time course of mean PFNa population activity, as measured via calcium signals, in the noduli as a function of airflow direction. The left nodulus is shown in blue, and the right nodulus is shown in red. The egocentric airflow direction is shown above each panel, and the time of airflow stimulus presentation is flanked by the dashed gray lines. **(C)** Tuning curve of PFNa-derived calcium signals in the noduli to egocentric airflow direction. Each point in this plot represents the mean value of the noduli z-score over the last 2 s of the airflow stimulus presentation in B. Note that the tuning curves for PFNa population activity in the noduli are double-phased. **(D)** Time course of mean LNOa population activity in the noduli as a function of airflow direction. The LNOa neurons are one of the numerically dominant synaptic inputs to the PFNa neurons in the noduli. The left nodulus is shown in blue, and the right nodulus is shown in red. The egocentric airflow direction is shown above each panel, and the time of airflow stimulus presentation is flanked by the dashed gray lines. **(E)** Tuning curve of LNOa-derived calcium signals in the noduli to egocentric airflow direction. Each point in this plot represents the mean value of the noduli z-score over the last 2 s of the airflow stimulus presentation in D. Note that the tuning curves for LNOa calcium have a single peak. **(F)** Cell-type specific expression of smFLAG-VGlut, a marker for glutamatergic neurons (strategy described in ref.^61^, and CD8-mCherry, a membrane marker, in the LNOa neurons. The driver line used was SS47432. Same as in A, but for cell-type specific expression of 9xV5-vGAT, a marker for GABAergic neurons^62^. **(G)** Comparison of the phase-nulled calcium signals of PFNa neurons in the absence of airflow (left panel) and when airflow is blown from directly behind the fly (180°, right panel). The horizontal dotted line marks the highest mean calcium responses in the no airflow condition, to aid comparison of the values in the two panels; note that both sinusoids peak at values above the line in the 180° airflow condition. **(H)** Reproduction of data from panels B and D (left two panels showing GCaMP signals), plus GFlamp1^75^ signals in PFNa neurons measured at the noduli to in response to airflow presented from the egocentric rearmost angle (180°). Note that whereas PFNa calcium shows minimal change during the stimulus presentation, the LNOa neurons are responsive. In addition, GFlamp1 signal rises within the PFNa neurons themselves shortly after the stimulus onset. **(I)** Sinusoidal function fits to the average phase-nulled PFNa calcium bumps elicited by frontal airflow (left panel) and airflow from the rear (right panel). The fit is shown as a dotted black line; the data is replotted from panel A, using the same color conventions. **(J)** Sinusoidal function fits to the average baseline-subtracted *Vm*, as a function of heading direction and airflow. The fit is shown as a dotted black line; the data is replotted from Figure 3C and 3E. Data from left-bridge-innervating PFNa neurons is shown in red, whereas data from right-bridge-innervating neurons is shown in blue.

**Figure S3.**
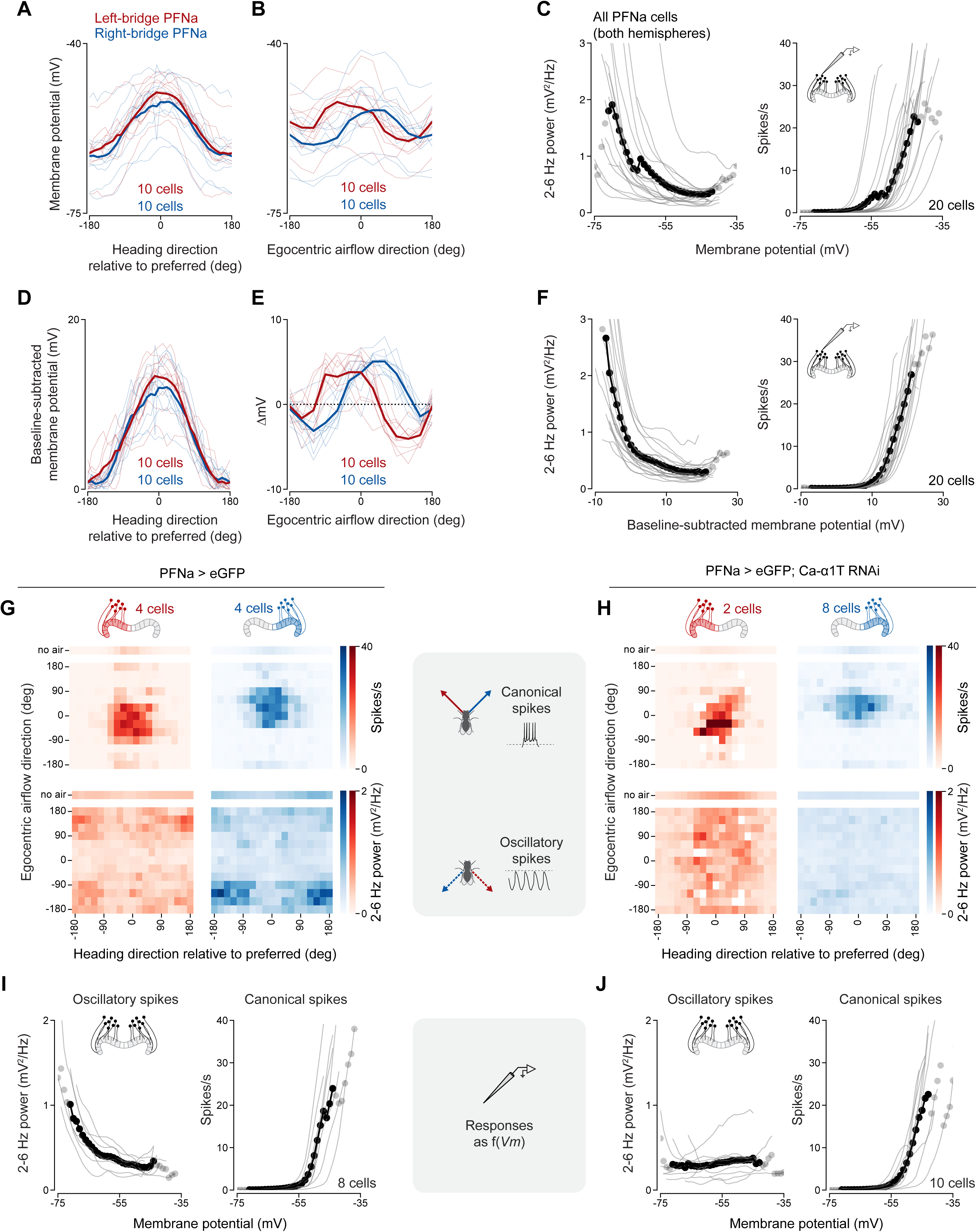
related to Figure 3. Raw and normalized *Vm* tuning curves from PFNa neurons. **(A)** Tuning of raw PFNa *Vm* to the fly’s heading, as estimated by the angular position of the closed-loop bar on the visual display. All tuning curves have been phase-aligned to have their peak at 0 (Methods). Left-bridge PFNa neurons: red. Right-bridge PFNa neurons: blue. Thin lines: single fly averages. Thick lines: population averages. Data from 20 cells (10 cells per hemisphere). **(B)** Raw *Vm* airflow direction tuning curves of PFNa neurons. These tuning curves have not been shifted. Thin lines: single fly averages. Thick lines: population averages. Data from 20 cells (10 cells per hemisphere). **(C)** 2-6 Hz power (left panel) and canonical spike rate (right panel) as a function of raw *Vm* in PFNa neurons. Responses of right- and left-bridge PFNa neurons were pooled. Thin lines: single fly averages. Thick lines: population averages. Mean points in which less than 50% of the curves contributed to the bin are displayed in gray. Data from 20 cells. **(D)** Same as in A, but showing the baseline-subtracted *Vm* instead of the raw *Vm*. The baseline was defined as the minimum value of the heading tuning curve (see Methods). **(E)** Same as in B, but showing the difference in *Vm* pre- and post-airflow stimulus instead of the raw *Vm*. **(F)** Same as in C, but showing the baseline-subtracted *Vm* instead of the raw *Vm*. The baseline was defined as the minimum value of the heading tuning curve (see Methods). Mean points in which less than 50% of the curves contributed to the bin are displayed in gray. **(G)** Conjunctive tuning of left- and right-bridge PFNa cells to the direction of airflow and heading. These recordings were made from flies of the genetic background used to create the TRiP RNAi libraries^30^ (“empty RNAi control”), but where eGFP was expressed in PFNa neurons. **(H)** Same as in panel G, but in PFNa cells carrying the construct TRiP.HMS01948, which allows for expression (under UAS control) of a double-stranded RNA that targets Ca-α1T transcripts for degradation (Ca-α1T RNAi). **(I)** Same as in C, but recording from PFNa cells in flies of the empty RNAi control genotype. **(J)** Same as in C, but recording from PFNa cells carrying the construct TRiP.HMS01948 (Ca-α 1T RNAi).

**Figure S4.**
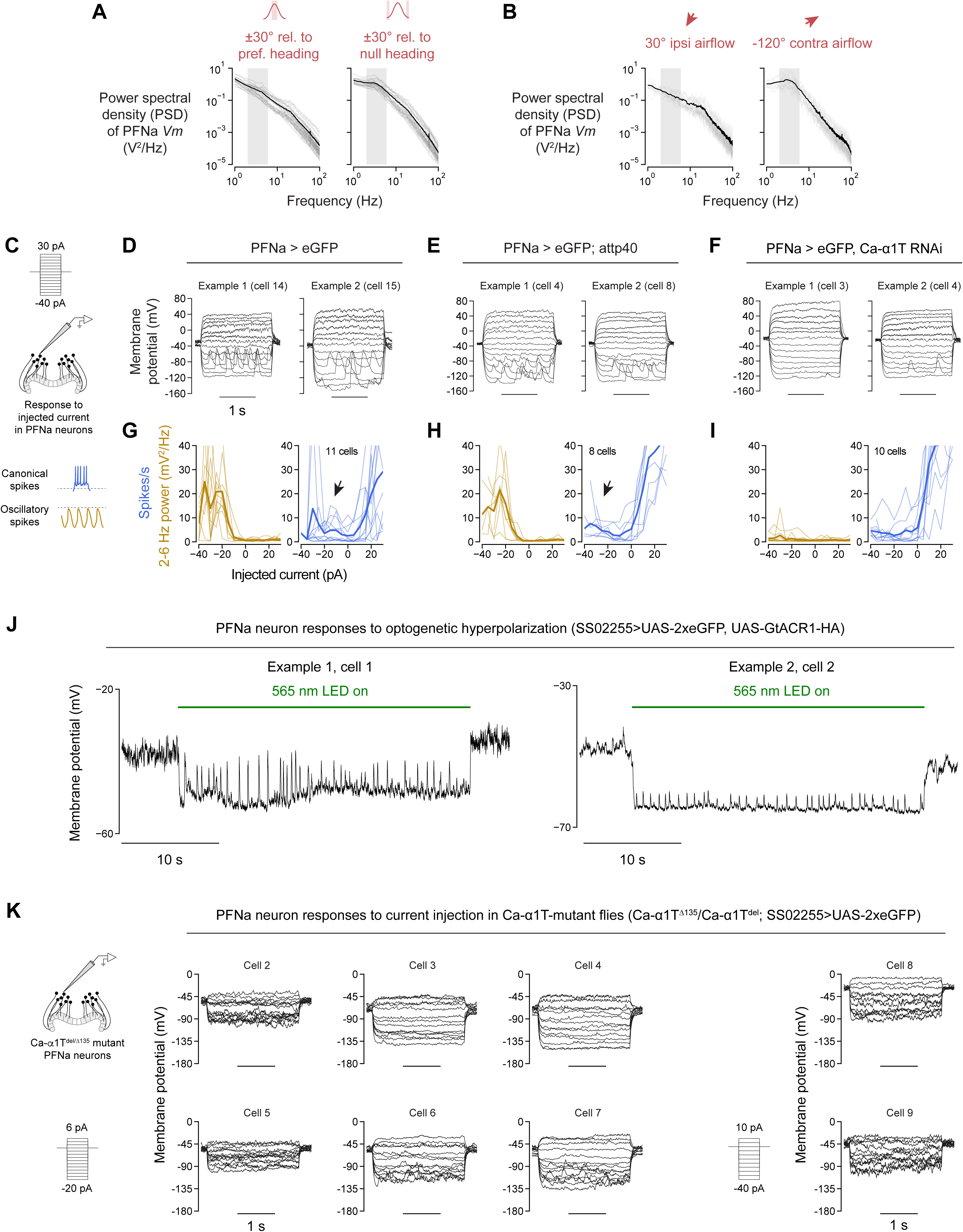
related to Figures 3 and 4. Features of the *Vm* of PFNa neurons, with and without knockdown of Ca-a1T channels. **(A)** Power spectral density (PSD) of the *Vm* of PFNa neurons during moments where the fly’s heading was aligned to the cell’s preferred direction (left panel) and moments where it was aligned to the null direction (right panel). Each panel shows a 60° bin of heading data (e.g. the left panel shows the PSD for the *Vm* of PFNa neurons when the fly’s heading was within ±30° relative to each cell’s preferred heading). The gray box indicates the 2-6 Hz band. Values from 20 cells. **(B)** Same as in A, but for depolarizing stimuli (airflow from 30° for right-bridge-innervating PFNa neurons and -30° for left-bridge-innervating PFNa neurons) and hyperpolarizing stimuli (airflow from -120° for right-bridge-innervating PFNa neurons and 120° for left-bridge-innervating PFNa neurons). Values from 20 cells. **(C)** Schematic of the current injection experiment. Each experiment consisted of a family of 15 current steps spaced 5 pA apart, starting at -40 pA and ending at 30 pA. We pooled cells innervating both brain hemispheres in this figure. **(D)** Representative families of current injections for 2 out of 11 different PFNa cells. This genotype is SS02255>UAS-2xeGFP, where no intentional perturbations were inserted. To the extent that it was possible, the recordings were preferentially acquired while the fly was quiescent. This was done to minimize depolarizing input from self-motion or membrane potential changes due to the visual stimulus entering or exiting the cell’s receptive field. **(E)** Same as in D, but in empty RNAi control genotype flies. **(F)** Same as in E, but in PFNa neurons expressing TRiP.HMS01948, which targets the Ca-α1T transcript (Ca-α1T RNAi). **(G)**2-6 Hz power (left panel, gold) and canonical spike responses (right panel, blue) as a function of injected current in PFNa neurons in SS02255>UAS-2xeGFP flies, where no intentional perturbations were inserted. The responses of right- and left-bridge innervating neurons were pooled. Thin lines represent individual cells, thick lines represent the average responses of 11 cells. The black arrow highlights a slight increase in the spike rate at negative current injections, it is due to sodium spikes that ride on top of calcium spikes. **(H)** Same as in G, but in empty RNAi control genotype flies. 8 cells. **(I)** Same as in (H), but in PFNa cells expressing 2xeGFP and TRiP.HMS01948, which targets the Ca-α1T transcript. 10 cells. **(J)** Two example traces of PFNa neurons showing calcium spikes in response to GtACR1-mediated hyperpolarization. We stimulated the fly using a 565 nm LED. The light intensity was measured to be 21 µW/mm2 prior to inserting a neutral density filter in the light path to reduce the magnitude of optogenetic stimulation. **(K)** Responses of Ca-α1T-mutant (Ca-α1Tdel/Δ135) PFNa neurons to current injection. The injection protocols are schematized on the bottom left of each set of cells. The total number of recorded cells is 8, data collected from 3 different flies.

**Figure S5.**
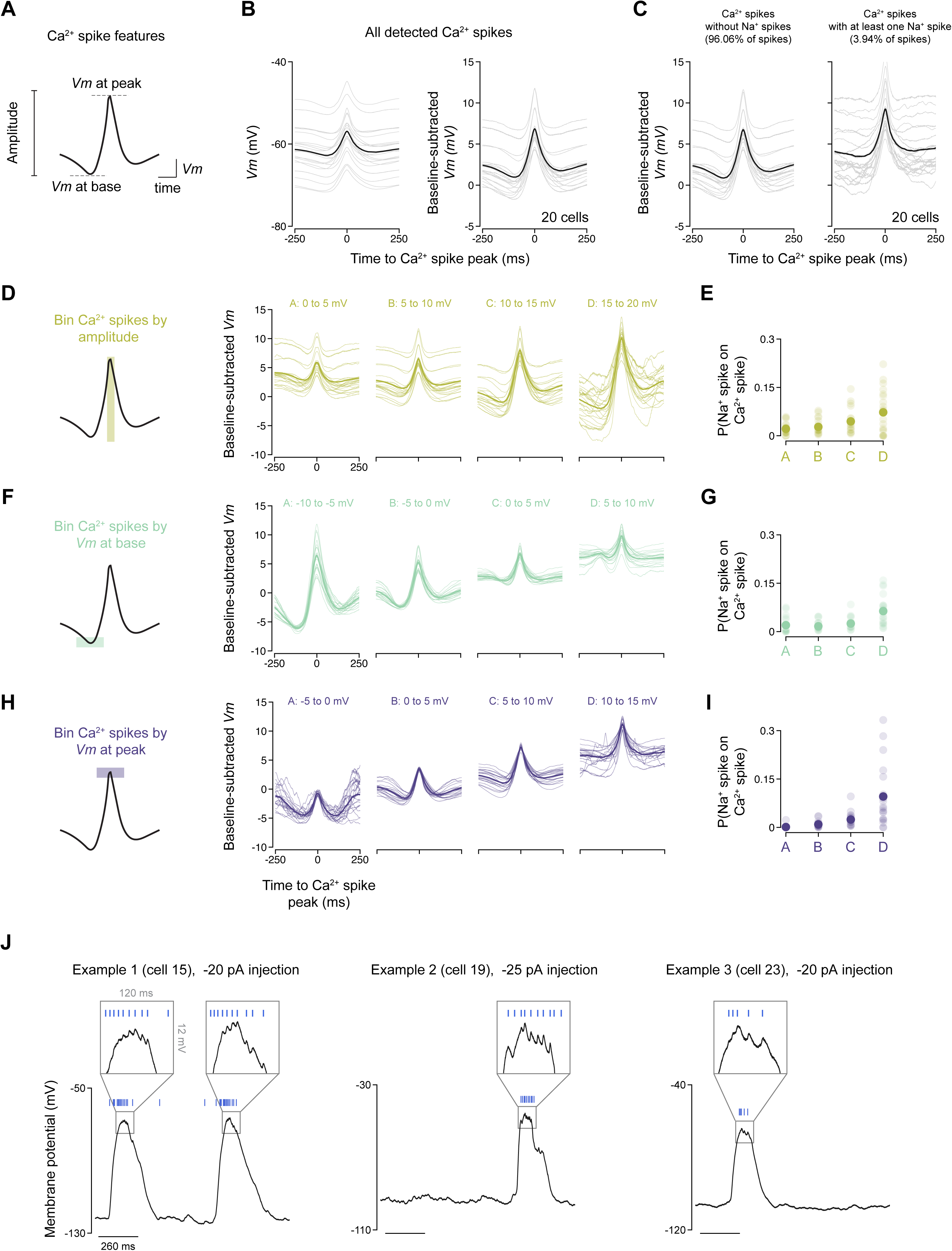
related to Figure 3. Characterization of the calcium spikes in PFNa neurons. **(A)** Schematic depicting features of individual calcium spikes. We used these features to bin the data in panels D through I. See Methods for details on calcium spike detection. **(B)** Average waveforms for all detected calcium spikes. The left panel’s y-axis shows the raw membrane potential, whereas the right panel’s y-axis displays the normalized membrane potential. The gray lines show the average waveform for individual cells, the black line shows the mean across all cells. Data from 68248 calcium spikes detected across 20 cells. The recordings were collected from 19 different flies. **(C)** Average waveforms for calcium spikes where we were not able to detect a sodium spike riding at the peak (left panel) and for calcium spikes where we did detect at least one sodium spike within 50 ms of the peak (right panel). The gray lines show the average waveform for calcium spikes detected in individual cells, the black line shows the mean across all cells. Data from 20 cells. **(D)** Average waveforms for calcium spikes binned by spike amplitude. The amplitude values used for each bin are displayed on top of each graph. Thin lines show average waveforms for one cell, thick lines show the mean across all cells. The four amplitude bins contain data from all cells (n=20). **(E)** Probability of a calcium spike displaying at least one sodium spike within 50 ms of the calcium spike peak as a function of calcium spike amplitude. We computed this probability simply by dividing the number of calcium spikes displaying sodium spikes by the total number of calcium spikes detected across the recording. Light-colored dots show values for individual cells, the dark-colored dots show the mean across cells. **(F)** Same as in D, but for calcium spikes binned by the *Vm* value preceding the spike. We define *Vm* at base as the average membrane potential between 150 and 100 ms preceding the calcium spike peak. **(G)** Same as in E, but for calcium spikes binned by *Vm* at the base of the spike. **(H)** Same as in D, but for calcium spikes binned by the value of *Vm* at the peak of the spike. In this case, condition “D” (spikes with Vm at peak within 10-15 mV) only shows data for 19 out of 20 cells. This is because one cell did not display spikes that fell into the 10-15 mV *Vm* at peak category. **(I)** Same as in E, but for calcium spikes binned by the value of *Vm* at the peak of the spike. **(J)** Three example traces of current injection steps that yielded sodium spikes at the peak of a calcium spike. The insets are magnified epochs highlighting the sodium spikes (blue lines). (Note that the cell bodies of *Drosophila* neurons connect to the cell arbors through a thin neurite that considerably filters voltage signals from the processes. Spikes in the PFNa neuron recordings are generally hard to visualize in raw voltage traces but are easily detected by filtering and thresholding the membrane potential.)

**Figure S6.**
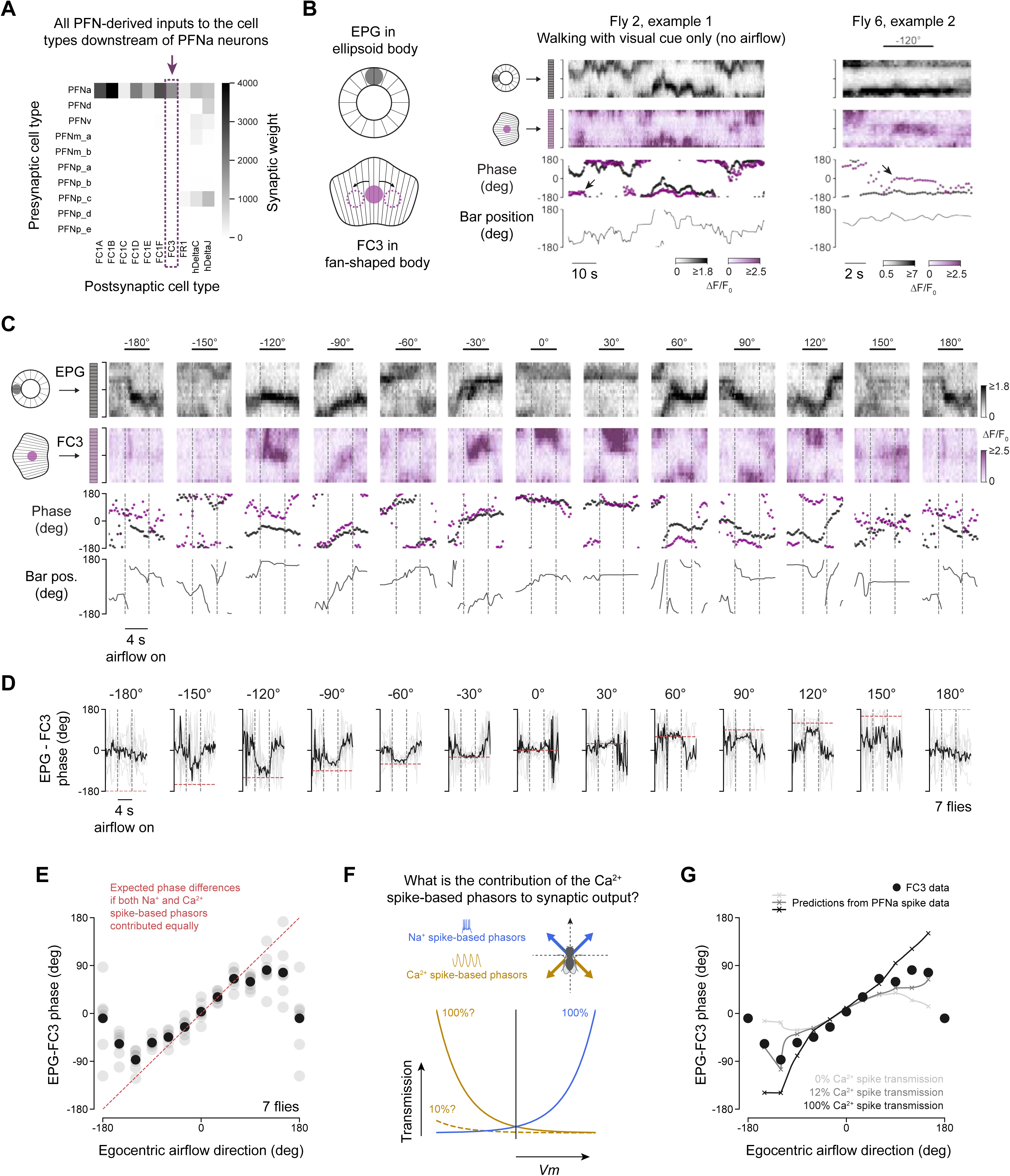
related to Figure 6. Assessment of the ability of FC3 neurons to signal the allocentric direction of air puffs. **(A)** Connectivity matrix showing all of the identified PFN neurons in the *Drosophila* hemibrain connectome^60^, and how they output onto the 10 types of columnar cell classes postsynaptic to PFNa neurons: the FC1A, FC1B, FC1C, FC1D, FC1E, FC1F, FC3, FR1, hΔC, and hΔJ cell classes^17^. The FR1, hΔC, and hΔJ neurons receive synaptic input from additional types of PFN cells, potentially introducing confounds into our analysis of the PFNa neurons’ synaptic output; we thus chose not to study these cell classes. The FC1 and FC3 neurons, in contrast, receive essentially all of their PFN-related synaptic input from PFNa neurons. The FC1 cells are the largest recipients of monosynaptic PFNa neuron input, but the six subtypes (FC1A-E) are challenging to parse using light microscopy. Therefore, we focused on the FC3 neurons, which have an identifiable anatomy, as our model postsynaptic neuron class. The FC3 neurons also receive heavy synaptic input from the FC1 neurons (not shown in this graph). **(B)** Example traces of two experiments where we simultaneously imaged the EPG and FC3 neurons. The left panel shows EPG and FC3 activity in the absence of airflow, whereas the right panel shows an example trace with an air puff from -120°. The EPG bump was imaged in the ellipsoid body, whereas the FC3 were imaged in the fan-shaped body, pictured here is the FC3 activity in layer 5 of the FB. The phase of the two population signals is overlaid in the third row, and the position of the visual stimulus on the screen is shown on the fourth row. The arrows highlight instances of a large difference between the phase of the FC3 neurons and the phase of the EPG neurons. **(C)** Example airflow responses of the EPG and FC3 neurons at the 12 airflow angles tested. The airflow-on period is flanked by the dotted lines, and the content and structure of the panels is otherwise the same as in B. **(D)** Time course of the difference of the FC3 phase position relative to the EPG phase during periods of airflow stimulation. The airflow-on period is flanked by the dotted lines, and the red dotted line denotes the expected magnitude of the phase difference if the PFNa system were operating as a four-vector set with equal amplitudes. The thin lines show individual flies, the thick lines show the average values across 7 flies. **(E)** Difference of the FC3 phase position relative to the EPG phase in the context of airflow stimulation like the one shown in D. The values shown here correspond to the average phase difference over the last two seconds of the airflow stimulus. The gray dots show individual flies, the black dots show the average values across 7 flies. The expected phase differences if the front and back vectors contributed equally are noted in red. **(F)** Schematic depicting how the sodium-spike-based phasors and calcium-spike-based phasors could contribute with different weights to synaptic transmission. The curves schematized here are drawn after the spike-rate and oscillation-power responses shown in Figure 7, but the y-axis shows the strength with which each type of signal might contribute to synaptic transmission from the PFNa neurons and therefore the calcium phase of the FC3 neurons. In this schematic, we assume the sodium-spike-based response contributes 100% to synaptic transmission, and ask whether the calcium spikes contribute just as much as the sodium spikes (gold curve labeled “100%”) or a smaller percentage (e.g. 10% as in the dashed gold curve). **(G)** Predicted phase of the postsynaptic FC3 neurons using the spike-rate and oscillation-power data from the PFNa neurons (from Figures 3H and 3I, see Methods). Each line shows the expected FC3 response given varying contributions of the calcium spikes to fast synaptic transmission assuming the vector axes in Figure S7F (±39° for the sodium-spike-vectors and ±124° for the calcium spike vectors). The light gray line shows the expected FC3 response if the sodium-spike-based PFNa phasors alone were to determine the FC3 phase. The black line shows the expected response if the sodium and calcium spikes were to contribute equally to synaptic transmission. The dark gray curve shows the best-fit percentage of calcium spike contribution, which was 12% for these physiologically-determined vector angles; the same result was obtained assuming ±45° or ±55° for the front angles. The black markers show the mean data from panel E. Data corresponding to airflow from 180° cannot be explained by this analysis and was thus excluded from the fits.

**Figure S7.**
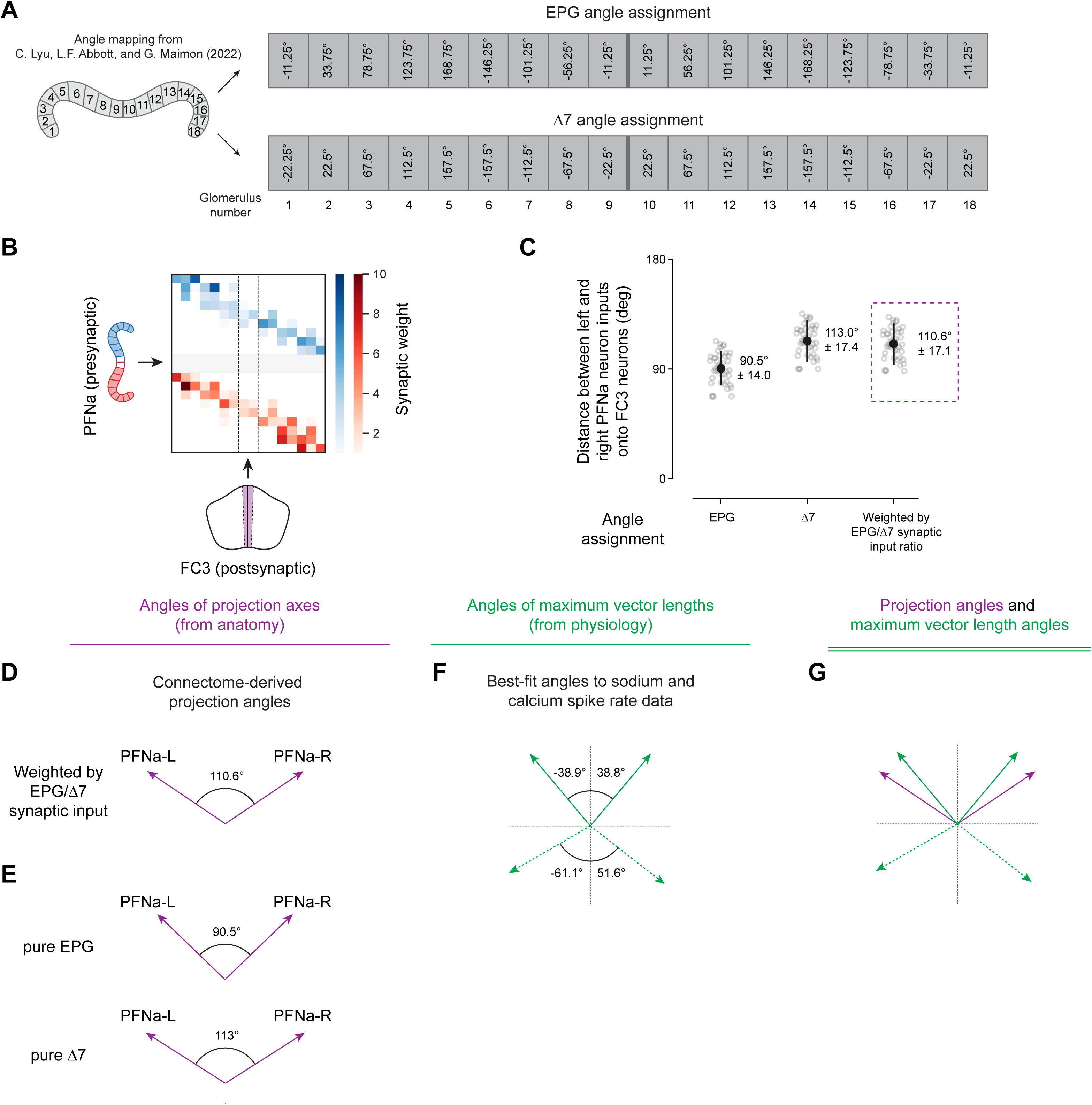
related to Figure 7. Analysis of the vector-axis directions and peak-vector length angles in PFNa neurons. **(A)** Angular indexing of the protocerebral bridge, with different schemes for the EPG neurons and the Δ7 neurons. These angles were experimentally defined in Lyu et al 202211 and we adopt them in this study. Adjacent glomeruli are generally spaced 45° apart in both cell types, but the indexing values of the Δ7 neurons are shifted to those from the EPG neurons by 1/4th of a glomerulus (11.25°). Note that the EPG neurons do not innervate the outermost two glomeruli (1 and 18); these angles were inferred given the bridge innervation pattern of the PEN^24,63^ (and PFN) neurons. **(B)** Connectivity matrix of the PFNa neurons onto the FC3 neurons, averaged over neuron instances that innervate the same bridge glomerulus or fan-shaped body column. Every fan-shaped body column innervated by the FC3 neurons receives input from both the left- and the right-bridge PFNa neurons. **(C)** Angle difference between left-bridge and right-bridge PFNa neuron inputs onto the FC3 neurons. Each circle corresponds to the values calculated for individual FC3 neurons, using the bridge indexing of either EPG alone, Δ7 alone, or a weighted combination. **(D)** Anatomically defined angle between the left and right PFNa vectors. 110.6° corresponds to the value calculated in C via connectome analysis. Because the front and rear vectors in the PFNa system arise from the same anatomical inputs, the pink arrows could point both forward or backward; we draw them pointing forward for simplicity. **(E)** Same as in D, but assuming pure EPG or pure Δ7 input. **(F)** We fit the heading-and-airflow 2-D heatmaps for spiking responses using the quadratic model in Figure 5. The best-fit angles for the preferred airflow directions are shown as the angles for each of the four vectors. **(G)** Overlay of the angles of the projection axes (purple) and the angles of the maximum vector lengths (green) as estimated in panels D and F.

**Table S1.** Experimental genotypes and cross details, related to Figures 2, 3, 4, 6, 7, S1, S2, S3, S4, S5, and S6.

| Figure | Parent 1 | Parent 2 | Experimental genotype |
| --- | --- | --- | --- |
| Figure 2B, 2C, 2D, 2E, 2F;<br>Figure 7F;<br>Figure S2A, S2G, S2I; | w+;<br>LexAop-RGECO1a in su(Hw)attp5;<br>UAS-jGCaMP7f in VK00005 | w+;<br>60D05-LexA in attp40;<br>30E10-Gal4 in attp2 | w+;<br>60D05-LexA/LexAop-RGECO1a; 30E10-Gal4/UAS-jGCaMP7f |
| Figure 2G;<br>Figure S2B, S2C, S2H | w+;<br>;<br>UAS-jGCaMP7f in VK00005 | 30E10-Gal4 in attp2 | w+; ;<br>30E10-Gal4/UAS-jGCaMP7f |
| Figure 3B, 3C, 3E, 3F, 3G, 3H, 3I;<br>Figure 7B, 7C, 7D;<br>Figure S2J;<br>Figure S3A, S3B, S3C, S3D, S3E, S3F,<br>Figure S4A, S4B, S4D, S4G;<br>Figure S5B, S5C, S5D, S5E, S5F, S5G, S5H, S5I, S5J; | w+;<br>UAS-2xeGFP (in Chr 2) | SS02255<br>(VT016D01-p65ADZp in attp40;<br>VT016114-ZpGdbd in attp2) | w+;<br>SS02255/UAS-2xeGFP |
| Figure 4C | Ca-α1T<sup>GFP</sup> | Ca-α1T<sup>GFP</sup> | Ca-α1T<sup>GFP</sup> |
| Figure 4D;<br>Figure S4E, S4H | w+;<br>attp40;<br>UAS-Dcr2 (in Chr 3) | w+;<br>UAS-2xeGFP (in Chr 2);<br>30E10-Gal4 in attp2 | w+;<br>attp40/UAS-2xeGFP;<br>30E10-Gal4/UAS-Dcr2 |
| Figure 4G;<br>Figure S4F; S4I | w+;<br>TRiP.HMS01948 in attp40;<br>UAS-Dcr2 (in Chr 3) | w+;<br>UAS-2xeGFP (in Chr 2);<br>30E10-Gal4 in attp2 | w+;<br>TRiP.HMS01948/UAS-2xeGFP;<br>30E10-Gal4/UAS-Dcr2 |
| Figure 4E, 4F | w+;<br>attp40;<br>UAS-Dcr2 (in Chr 3) | 30E10-Gal4 in attp2,<br>UAS-jGCaMP7f in VK00005 | w+/w;<br>attp40/+;<br>30E10-Gal4, UAS-jGCaMP7f/UAS-Dcr2 |
| Figure 4H, 4I | w+;<br>TRiP.HMS01948 in attp40 | 30E10-Gal4 in attp2,<br>UAS-jGCaMP7f in VK00005 | w+/w;<br>TRiP.HMS01948/+;<br>30E10-Gal4/UAS-jGCaMP7f |
| Figure 6C | w+;<br>27F02-LexA in attp40,<br>LexAop-syt-jGCaMP8s in VK00022 | w+;<br>27F02-LexA in attp40,<br>LexAop-syt-jGCaMP8s in VK00022 | w+;<br>27F02-LexA in attp40,<br>LexAop-syt-jGCaMP8s in VK00022 |
| Figure 6C | UAS-IVS-mCD8::GFP in attp2 | 12E04-LexA in attp40 | w;<br>12E04-LexA;<br>UAS-IVS-mCG8::GFP |
| Figure 6E, 6F, 6G, 6H | w+;<br>27F02-LexA in attp40,<br>LexAop-syt-jGCaMP8s in VK00022;<br>UAS-CsChrimson-tdTomato in attp2 | w+;<br>12E04-LexA in attp40;<br>41H07-Gal4 in attp2 | w+;<br>27F02-LexA, LexAop-syt-jGCaMP8s/12E04-LexA;<br>41H07-Gal4/UAS-CsChrimson-tdTomato |
| Figure 6F, 6G, 6H | w<sup>+</sup>;<br>27F02-LexA in attp40,<br>LexAop-syt-jGCaMP8s in<br>VK00022;<br>UAS-CsChrimson-tdTomato in<br>attp2 | w<sup>+</sup>;<br>12E04-LexA in attp40;<br>VT056655-Gal4 in attp2 | w<sup>+</sup>;<br>27F02-LexA, LexAop-syt-<br>jGCaMP8s/12E04-LexA;<br>VT056655-Gal4/UAS-<br>CsChrimson-tdTomato |
| Figure 6F, 6G, 6H | w<sup>+</sup>;<br>27F02-LexA in attp40,<br>LexAop-syt-jGCaMP8s in<br>VK00022;<br>UAS-CsChrimson-tdTomato in<br>attp2 | w<sup>+</sup>;<br>12E04-LexA in attp40;<br>empty Gal4 in attp2 | w<sup>+</sup>;<br>27F02-LexA, LexAop-syt-<br>jGCaMP8s/12E04-LexA;<br>empty Gal4/UAS-<br>CsChrimson-tdTomato |
| Figure 6J, 6K, 6L,<br>6M | w<sup>+</sup>;<br>27F02-LexA in attp40,<br>LexAop-syt-jGCaMP8s in<br>VK00022;<br>UAS-GtACR-HA in VK00005 | w<sup>+</sup>;<br>12E04-LexA in attp40;<br>41H07-Gal4 in attp2 | w<sup>+</sup>;<br>27F02-LexA, LexAop-syt-<br>jGCaMP8s/12E04-LexA;<br>41H07-Gal4/UAS-GtACR-<br>HA |
| Figure 6K, 6L, 6M | w<sup>+</sup>;<br>27F02-LexA in attp40,<br>LexAop-syt-jGCaMP8s in<br>VK00022;<br>UAS-GtACR-HA in VK00005 | w<sup>+</sup>;<br>12E04-LexA in attp40;<br>VT056655-Gal4 in attp2 | w<sup>+</sup>;<br>27F02-LexA, LexAop-syt-<br>jGCaMP8s/12E04-LexA;<br>VT056655-Gal4/UAS-<br>GtACR-HA |
| Figure 6K, 6L, 6M | w<sup>+</sup>;<br>27F02-LexA in attp40,<br>LexAop-syt-jGCaMP8s in<br>VK00022;<br>UAS-GtACR-HA in VK00005 | w<sup>+</sup>;<br>12E04-LexA in attp40;<br>empty Gal4 in attp2 | w<sup>+</sup>;<br>27F02-LexA, LexAop-syt-<br>jGCaMP8s/12E04-LexA;<br>empty Gal4/UAS-GtACR-<br>HA |
| Figure 6O, 6P,<br>6Q, 6R | Ca-α1T<sup>del</sup>;<br>27F02-LexA in attp40,<br>LexAop-syt-jGCaMP8s in<br>VK00022;<br>UAS-GtACR-HA in VK00005 | Ca-α1T<sup>Δ135</sup>;<br>12E04-LexA in attp40;<br>41H07-Gal4 in attp2 | Ca-α1T<sup>del</sup>/Ca-α1T<sup>Δ135</sup>;<br>27F02-LexA, LexAop-syt-jGCaMP8s/12E04-LexA;<br>41H07-Gal4/UAS-GtACR-HA |
| Figure 6P, 6Q, 6R | Ca-α1T<sup>del</sup>;<br>27F02-LexA in attp40,<br>LexAop-syt-jGCaMP8s in<br>VK00022;<br>UAS-GtACR-HA in VK00005 | w<sup>+</sup>;<br>12E04-LexA in attp40;<br>41H07-Gal4 in attp2 | Ca-α1T<sup>del</sup>/w<sup>+</sup>;<br>27F02-LexA, LexAop-syt-<br>jGCaMP8s/12E04-LexA;<br>41H07-Gal4/UAS-GtACR-<br>HA |
| Figure S1A, S1B,<br>S1C, S1D, S1E,<br>S1F, S1G, S1H,<br>S1I;<br>Figure S6B, S6C,<br>S6D, S6E, S6G | LexAop-jGCaMP7f in<br>su(Hw)attp8;<br>;<br>UAS-jRGECO1a in VK00005 | w<sup>+</sup>;<br>12E04-LexA in attp40;<br>60D05-Gal4 in attp2 | LexAop-jGCaMP7f/w<sup>+</sup>;<br>12E04-LexA;<br>60D05-Gal4/UAS-<br>jRGECO1a |
| Figure S2D, S2E,<br>S2H | w<sup>+</sup>;<br>;<br>UAS-jGCaMP7f in VK00005 | SS47432<br>(VT046049-p65ADZp in<br>attp40;<br>VT024603-ZpGdbd in<br>attp2) | w<sup>+</sup>;<br>SS47432/UAS-jGCaMP7f |
| Figure S2F | w;<br>VGlut1[B2RT.STOP.smFLAG],<br>20XUAS-DSCP-B2R in<br>VK00018;<br>UAS-mCD8.mCherry | w<sup>+</sup>;<br>SS47432<br>(VT046049-p65ADZp in<br>attp40;<br>VT024603-ZpGdbd in<br>attp2) | w<sup>+</sup>/w;<br>SS47432/ B2RT- STOP-<br>B2RT-smFLAG-vGlut,<br>UAS-mCD8.mCherry |
| Figure S2F | w;<br>VGAT[RSRT(STOP).9XV5],<br>20XUAS-RSR.S in JK22C;<br>UAS-mCD8.mCherry | w<sup>+</sup>;<br>SS47432<br>(VT046049-p65ADZp in<br>attp40;<br>VT024603-ZpGdbd in<br>attp2) | w<sup>+</sup>/w;<br>SS47432/<br>RSRT-STOP-RSRT-<br>9XV5-vGAT, UAS-<br>mCD8.mCherry |
| Figure S2H | UAS-GFlamp1 in VK00005 | w<sup>+</sup>;<br>SS02255<br>(VT016D01-p65ADZp in<br>attp40;<br>VT016114-ZpGdbd in<br>attp2) | w<sup>+</sup>/yw;<br>SS02255/UAS-GFlamp1 |
| Figure S4J | w<sup>+</sup>;<br>UAS-2xeGFP;<br>UAS-GtACR-HA in VK00005 | w<sup>+</sup>;<br>SS02255<br>(VT016D01-p65ADZp in<br>attp40;<br>VT016114-ZpGdbd in<br>attp2) | w<sup>+</sup>;<br>SS02255/UAS-2xeGFP;<br>UAS-GtACR-HA |
| Figure S4K | Ca-α1T<sup>del</sup>;<br>UAS-2xeGFP (in Chr 2) | Ca-α1T<sup>Δ135</sup>;<br>SS02255<br>(VT016D01-p65ADZp in<br>attp40;<br>VT016114-ZpGdbd in<br>attp2) | Ca-α1T<sup>del</sup>/Ca-α1T<sup>Δ135</sup>;<br>SS02255/UAS-2xeGFP |

**Table S2.** Passive membrane properties of the patch-clamp recordings in this study, related to Figures 3, 4, 7, S3, S4, and S5.

| Dataset | Fly ID | Cell ID | Recording ID | Cm (pF) | Rm (GOhm) | Ra (MOhm) |
| --- | --- | --- | --- | --- | --- | --- |
| wild-type | A | 5 | 2022_01_17_0002 | 2.39 | 2.3 | 31.8 |
| wild-type | B | 6 | 2022_01_18_0002 |  |  |  |
| wild-type | C | 7 | 2022_01_19_0002 | 5.13 | 2.3 | 9.8 |
| wild-type | D | 9 | 2022_01_20_0014 |  |  |  |
| wild-type | E | 10 | 2022_01_21_0000 | 3.12 | 1.5 | 16.2 |
| wild-type | E | 11 | 2022_01_21_0006 |  |  |  |
| wild-type | F | 12 | 2022_01_25_0006 |  |  |  |
| wild-type | G | 13 | 2022_01_26_0000 | 1.92 | 1.2 | 26.6 |
| wild-type | H | 14 | 2022_01_27_0000 |  |  |  |
| wild-type | I | 15 | 2022_01_28_0000 |  |  |  |
| wild-type | J | 16 | 2022_02_03_0006 |  |  |  |
| wild-type | K | 17 | 2022_04_19_0001 | 0.941 | 1.6 | 55 |
| wild-type | L | 18 | 2022_04_19_0006 | 2.18 | 2 | 23.2 |
| wild-type | M | 20 | 2022_04_29_0003 | 2.7 | 1.3 | 18.7 |
| wild-type | N | 21 | 2022_05_02_0002 | 1.73 | 1.5 | 29.4 |
| wild-type | O | 22 | 2022_05_02_0008 | 2.41 | 2.7 | 20.9 |
| wild-type | P | 24 | 2022_05_04_0004 | 2.85 | 1.5 | 17.8 |
| wild-type | Q | 25 | 2022_05_05_0003 | 1.43 | 1.8 | 35.6 |
| wild-type | R | 26 | 2022_05_05_0008 | 1.82 | 2 | 27.8 |
| wild-type | S | 27 | 2022_05_31_0004 | 1.41 | 2.3 | 35.9 |
| Ca-a1T-RNAi | A | 1 | 2022_04_11_0001 | 2.5 | 1.6 | 20.3 |
| Ca-a1T-RNAi | B | 2 | 2022_04_11_0007 | 8.43 | 1.5 | 24.9 |
| Ca-a1T-RNAi | C | 3 | 2022_04_13_0001 | 3 | 2.3 | 16.8 |
| Ca-a1T-RNAi | C | 4 | 2022_04_13_0006 | 2.44 | 1.8 | 20.7 |
| Ca-a1T-RNAi | D | 5 | 2022_04_15_0001 | 1.61 | 2.3 | 31.5 |
| Ca-a1T-RNAi | D | 6 | 2022_04_15_0007 |  |  |  |
| Ca-a1T-RNAi | E | 7 | 2022_04_18_0002 | 1.41 | 1.5 | 36.2 |
| Ca-a1T-RNAi | E | 8 | 2022_04_18_0006 | 3.69 | 1.8 | 13.7 |
| Ca-a1T-RNAi | F | 9 | 2022_04_21_0002 | 1.92 | 2 | 26.4 |
| Ca-a1T-RNAi | G | 10 | 2022_04_25_0001 | 2.78 | 1.8 | 18.2 |
| empty-RNAi | A | 1 | 2022_06_15_0001 | 2.66 | 1.2 | 19.1 |
| empty-RNAi | A | 2 | 2022_06_15_0003 | 4.34 | 1.5 | 354.6 |
| empty-RNAi | B | 3 | 2022_06_15_0005 |  |  |  |
| empty-RNAi | C | 4 | 2022_06_16_0002 |  |  |  |
| empty-RNAi | D | 5 | 2022_06_20_0000 |  |  |  |
| empty-RNAi | E | 6 | 2022_06_20_0003 |  |  |  |
| empty-RNAi | F | 7 | 2022_06_21_0001 | 2.57 | 2.3 | 19.6 |
| empty-RNAi | F | 8 | 2022_06_21_0003 |  |  |  |

**Note:** We calculated Rm and Cm values using an online membrane test (in pClamp) that relies on fitting current response to voltage pulses in voltage clamp mode. Missing entries in this table are instances where either the fitting procedure did not yield stable Rm/Cm values, or where results of the membrane test were not recorded. The quality of recordings as assessed by baseline $V_m$ , spike rate, and $\text{Na}^+/\text{Ca}^{2+}$ spike thresholds (in current clamp mode) are comparable for cells with and without reported membrane test results.

**Table S3.** Statistical testing, related to Figures 4, 6, S1, and S4.

| Figure | Groups compared | Test | Null hypothesis | P-value<br>(**<u>P<0.05</u>**<br>**in bold**) |
| --- | --- | --- | --- | --- |
| 4D, 4G | Mean 2-6 Hz power below -60 mV for attp40 (*panel 4D, left side*) vs Ca-α1T RNAi (*panel 4G, left side*) | t-test, two-sided, means of two independent samples | Samples have equal means | **1.13x10<sup>-3</sup>** |
| 4F, 4I | Absolute R-L noduli z score at -120° airflow for attp40 (*panel 4F, left side*) vs Ca-α1T RNAi (*panel 4I, left side*) | t-test, two-sided, means of two independent samples | Samples have equal means | **1.13x10<sup>-2</sup>** |
| 4F, 4I | Absolute R/L noduli z score at 120° airflow for attp40 (*panel 3F, left side*) vs Ca-α1T RNAi (*panel 3G, left side*) | t-test, two-sided, means of two independent samples | Samples have equal means | **1.62x10<sup>-2</sup>** |
| 6H | Empty-Gal4 vs 41H07-Gal4 | t-test, two-sided, means of two independent samples | Samples have equal means | **4.27x10<sup>-4</sup>** |
| 6H | Empty-Gal4 vs VT056655-Gal4 | t-test, two-sided, means of two independent samples | Samples have equal means | **3.68x10<sup>-2</sup>** |
| 6M | Empty-Gal4 vs 41H07-Gal4 | Watson-Williams test for equality of the means of two or more samples of circular data | Samples have equal means | **4.83x10<sup>-9</sup>** |
| 6M | Empty-Gal4 vs VT056655-Gal4 | Watson-Williams test for equality of the means of two or more samples of circular data | Samples have equal means | **9.16x10<sup>-9</sup>** |
| 6R | Heterozygous (+/del) vs null (del/Δ135) | Watson-Williams test for equality of the means of two or more samples of circular data | Samples have equal means | **6.89x10<sup>-9</sup>** |
| S1E | No airflow vs airflow | t-test, two-sided, means of two paired samples | Samples have equal means | 0.725 |
| S1G | No airflow vs airflow | t-test, two-sided, means of two paired samples | Samples have equal means | 0.906 |
| S1H | No airflow vs airflow | t-test, two-sided, means of two paired samples | Samples have equal means | 0.290 |
| S4H, S4I | Mean 2-6 Hz power for current injections below -20 pA for attp40 (*panel S4H, left side*) vs Ca-α1T RNAi (*panel S4I, left side*) | t-test, two-sided, means of two independent samples | Samples have equal means | **1.91x10<sup>-5</sup>** |

## Key resources table

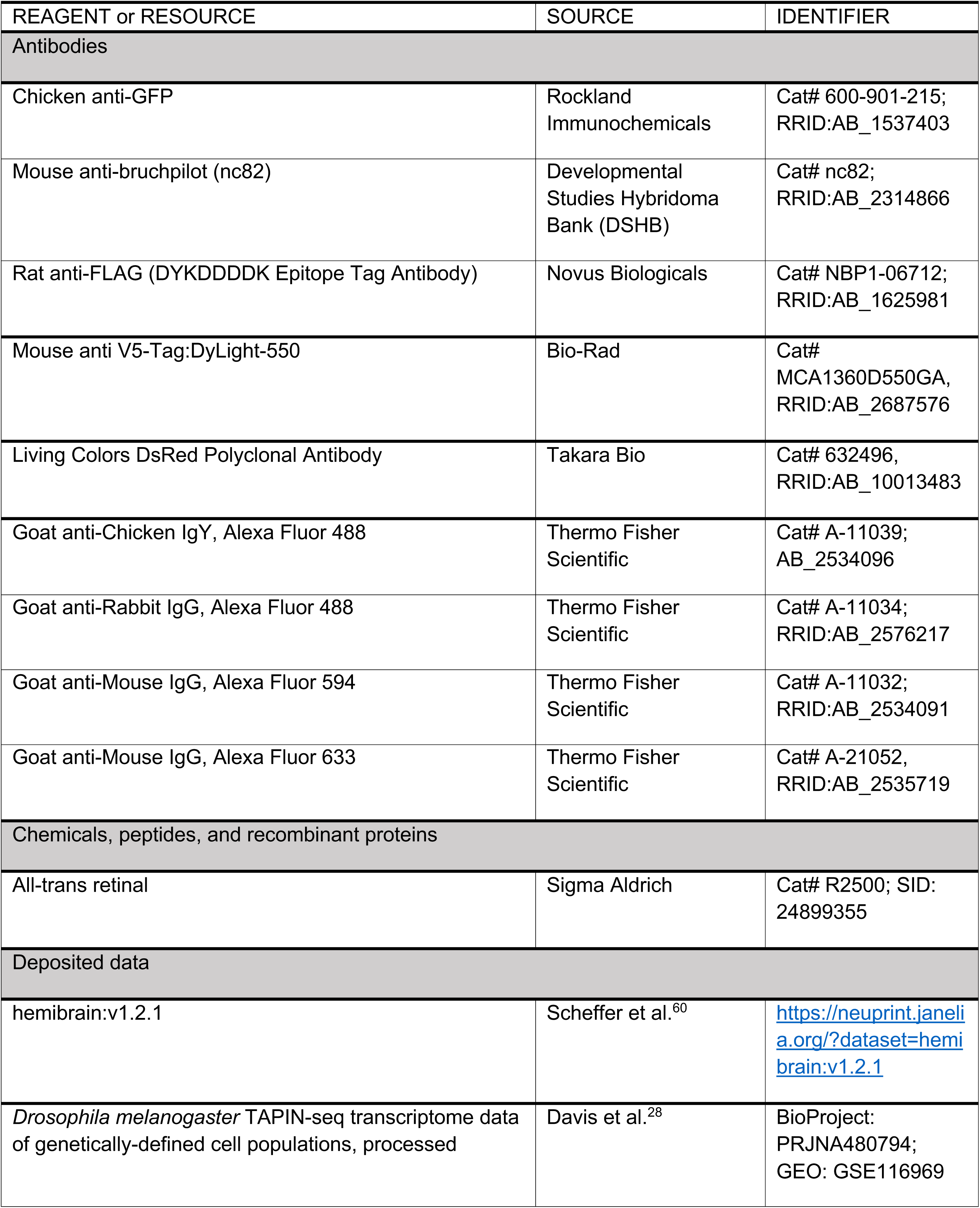
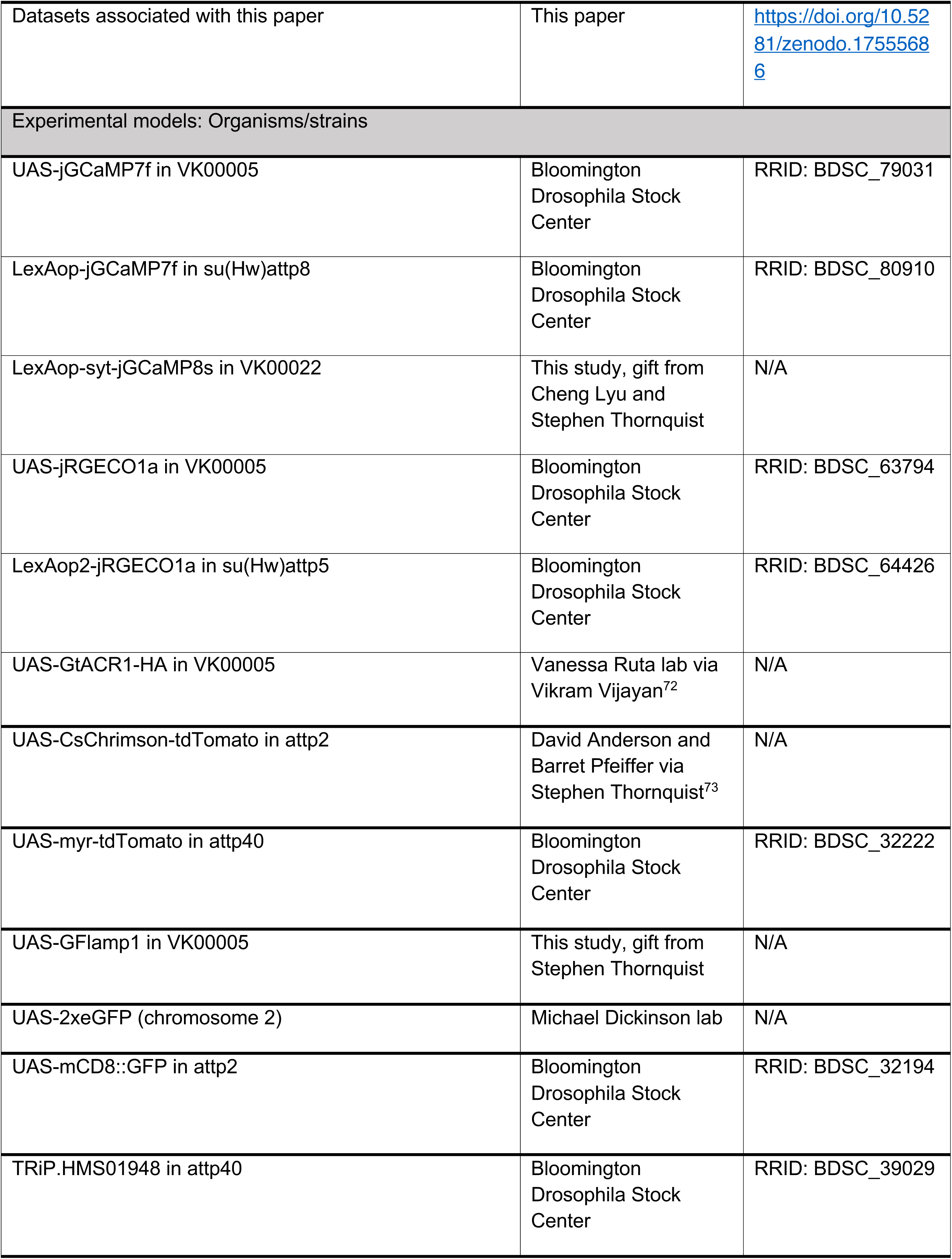
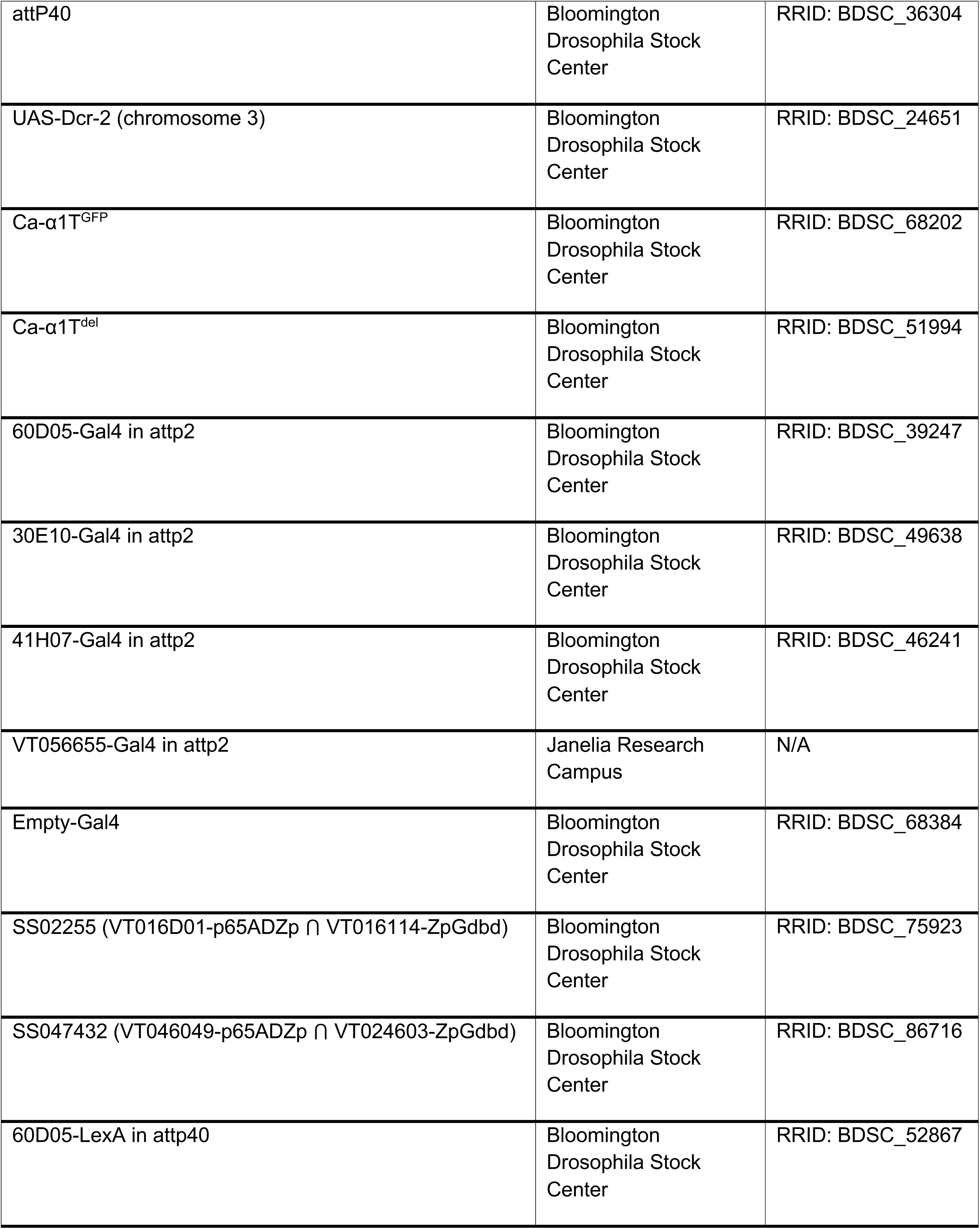
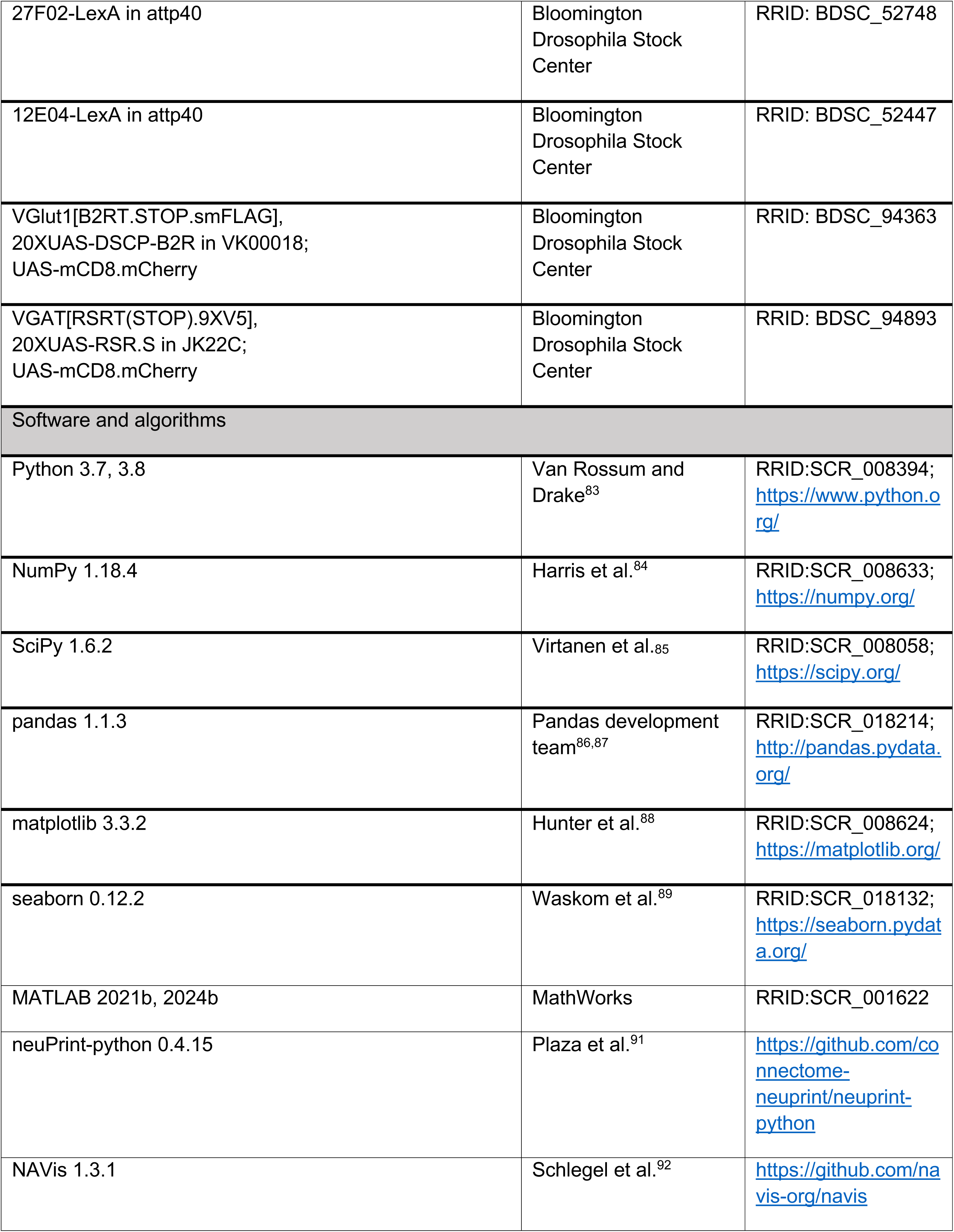
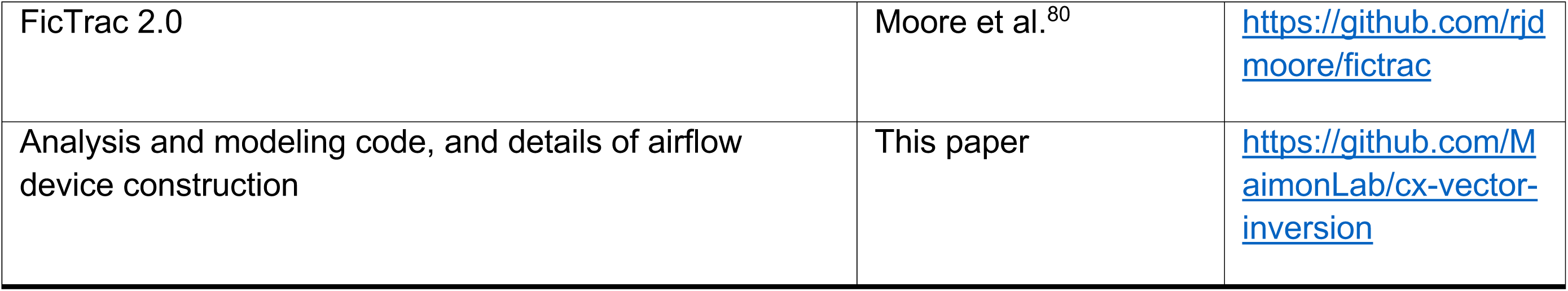

